# Distinct Mechanisms of Innate and Adaptive Immune Regulation Underlie Poor Oncologic Outcomes Associated with *KRAS-TP53* Co-Alteration in Pancreatic Cancer

**DOI:** 10.1101/2022.05.01.490244

**Authors:** Jashodeep Datta, Anna Bianchi, Iago De Castro Silva, Nilesh U. Deshpande, Long Long Cao, Siddharth Mehra, Samara Singh, Christine Rafie, Xiaodian Sun, Xi Chen, Xizi Dai, Antonio Colaprico, Prateek Sharma, Austin R. Dosch, Asha Pillai, Peter J. Hosein, Nagaraj S. Nagathihalli, Krishna V. Komanduri, Julie M. Wilson, Yuguang Ban, Nipun B. Merchant

**Affiliations:** Division of Surgical Oncology, Department of Surgery, University of Miami Miller School of Medicine, Miami, Florida, USA; Sylvester Comprehensive Cancer Center, Miami, Florida, USA; Biostatistics and Bioinformatics Shared Resource, Department of Public Health Sciences, University of Miami Miller School of Medicine, Miami, Florida, USA; Department of Surgery, University of Nebraska, Omaha, Nebraska, USA; Department of Pediatrics, University of Miami Miller School of Medicine, Miami, Florida, USA; Department of Microbiology and Immunology, University of Miami Miller School of Medicine, Miami, Florida, USA; Department of Medicine, University of Miami Miller School of Medicine, Miami, Florida, USA; PanCuRx Translational Research Initiative, Ontario, Institute for Cancer Research, Toronto, Ontario, Canada

## Abstract

Co-occurrent *KRAS* and *TP53* mutations define a majority of patients with pancreatic ductal adenocarcinoma (PDAC) and define its pro-metastatic proclivity. Here, we demonstrate that *KRAS*-*TP53* co-alteration is associated with worse survival compared with either *KRAS*-alone or *TP53*-alone altered PDAC in 245 patients with metastatic disease treated at a tertiary referral cancer center, and validate this observation in two independent molecularly annotated datasets. Compared with non-*TP53* mutated *KRAS*-altered tumors, *KRAS*-*TP53* co-alteration engenders disproportionately innate immune-enriched and CD8^+^ T-cell-excluded immune signatures. Leveraging *in silico, in vitro*, and *in vivo* models of human and murine PDAC, we discover a novel intersection between *KRAS*-*TP53* co-altered transcriptomes, *TP63*-defined squamous trans-differentiation, and myeloid-cell migration into the tumor microenvironment. Comparison of single-cell transcriptomes between *KRAS*-*TP53* co-altered and *KRAS*-altered/*TP53*^WT^ tumors revealed cancer cell-autonomous transcriptional programs that orchestrate innate immune trafficking and function. Moreover, we uncover granulocyte-derived inflammasome activation and *TNF* signaling as putative paracrine mediators of innate immunoregulatory transcriptional programs in *KRAS*-*TP53* co-altered PDAC. Immune subtyping of *KRAS*-*TP53* co-altered PDAC reveals conflation of intratumor heterogeneity with progenitor-like stemness properties. Coalescing these distinct molecular characteristics into a *KRAS*-*TP53* co-altered “immunoregulatory program” predicts chemoresistance in metastatic PDAC patients enrolled in the COMPASS trial, as well as worse overall survival.

## INTRODUCTION

Pancreatic ductal adenocarcinoma (PDAC) remains a tremendous clinical challenge due to its intrinsic and acquired therapeutic resistance. Although next-generation sequencing (NGS) has allowed comprehensive characterization of the dominant drivers of PDAC tumorigenesis— namely mutant *KRAS, TP53, SMAD4, CDKN2A*—these molecular alterations are not currently therapeutically actionable [1]. Beyond this “undruggable” genomic landscape in PDAC, defining hallmarks are its characteristic stromal desmoplasia as well as myeloid cell-enriched and T-cell-excluded tumor microenvironment (TME) [2]. However, how these genomic alterations that define PDAC tumorigenesis also dictate its immunosuppressive tumor biology and govern therapeutic resistance remains poorly understood, exposing a critically underexplored genotype-phenotype chasm.

To address the therapeutic impasse associated with the genotype-phenotype chasm in gastrointestinal (GI) tumors, our group has previously proposed an extreme-outlier model to identify cooperative genomic alterations that dictate prognosis and define therapeutic sensitivity or resistance [3-5]. This model also proposes comprehensive investigation of downstream transcriptomic, immunomic, and metabolic cellular consequences to reveal novel molecular vulnerabilities in therapeutically resistant tumors with high-risk genomic features [6]. Leveraging this methodology, our group has focused on the genomic co-occurrence of *KRAS* and *TP53* alterations as a prototype of this paradigm. Utilizing this extreme-outlier methodology, we have identified the association between *KRAS-TP53* co-alteration and a clinical phenotype defined by chemoresistance [3], aggressive non-salvageable metastatic proclivity [4, 5], and worse cancer-related survival compared with *KRAS* or *TP53* alterations alone [4] in *non-PDAC* GI cancers. Moreover, we and others have linked *KRAS-TP53* co-alteration with the disproportionate enrichment of innate immune populations and immunoregulatory signaling in these GI cancers, implicating *KRAS-TP53* cooperativity a model to investigate innate immune regulation [7].

Importantly, *KRAS-TP53* co-alteration is a foundational molecular event in PDAC tumorigenesis, modulating cellular signaling to induce the spontaneous development of PDAC in the well-established *LSL-K-ras*^*G12D/+*^; *Trp53*^*R172H/+*^; *Pdx-1*^*Cre/+*^ (KPC) genetically engineered mouse model (GEMM) [8]. Beyond sufficiency to induce spontaneous tumorigenesis, *Kras-Trp53* co-alteration also promotes invasion and generates a highly metastatic phenotype *in-vivo* [9]. Recent insight into the molecular underpinnings of the *KRAS-TP53* cooperativity-induced pro-metastatic phenotype revealed that oncogenic KRAS effectors activate CREB1 to allow physical interactions with mutant p53 that hyperactivate multiple pro-metastatic transcriptional networks [10]. In parallel, another recent study demonstrated the association of *TP53* missense mutations with high stromal cellularity and T-cell exclusion in PDAC [11]. However, the transcriptional programs differentially encoded by *KRAS-TP53* co-occurrence that govern innate immunoregulation and adaptive immune exclusion, as well as the association of such programs with therapeutic resistance and survival, have not been previously explored in PDAC.

In the present manuscript, we demonstrate that *KRAS-TP53* co-alteration in PDAC is not only associated with worse survival compared with *KRAS*-altered or *TP53* -altered PDAC in three independent datasets, but also engenders a disproportionately innate immune-enriched and CD8^+^ T-cell-excluded immune contexture. Leveraging several *in-silico, in-vitro*, and *in-vivo* models of human and murine PDAC, we discover a novel intersection between *KRAS-TP53* co-altered transcriptomes, *TP63*-defined squamous trans-differentiation, and myeloid cell migration and infiltration into the PDAC TME. Beyond cancer cell-autonomous transcriptional programs that orchestrate innate immune trafficking, we also uncover granulocyte-derived inflammasome activation and *TNF* signaling as putative paracrine mediators of immunoregulation in *KRAS-TP53* co-altered PDAC using single-cell RNA sequencing (scRNAseq) data. The CD8^+^ T-cell exclusion observed in *KRAS-TP53* co-altered PDAC appears to be driven by limited repertoire of molecular determinants reflective of adaptive immune dysfunction and immune evasion.

Consensus immune subtyping of *KRAS-TP53* co-altered PDAC reveals enrichment of intratumor heterogeneity and progenitor-like stemness properties—both associated with immune exclusion. Combining these novel molecular characteristics differentially overexpressed in *KRAS-TP53* co-altered PDAC reveals an immunoregulatory program (IRP) associated with chemotherapy resistance and worse overall survival in PDAC patients.

## METHODS

### UMiami cohort and independent validation datasets

After obtaining Institutional Review Board approval for the study, patients with metastatic PDAC who underwent treatment at the University of Miami (UMiami) between 7/2015 and 12/2019 and had available NGS testing using two commercially available platforms (Caris™ and FoundationOne™) were identified from PatientAtlas™, a repository harmonizing health system-wide NGS testing. Mutation calling was harmonized using the MC3 algorithm [12]. Patients with microsatellite-stable tumors and alterations in *KRAS, TP53*, or both (n=245) were retrieved and annotated by OncoKB knowledgebase [13] to designate putative driver or structural alterations. Oncoprint was generated utilizing a publicly available tool (https://www.cbioportal.org/oncoprinter). Clinical annotation of demographic and treatment-related variables were performed by retrospective review of the electronic medical record (**Table S1**). For survival analysis, patients were stratified into *KRAS-TP53* co-altered (n=171), *KRAS*-altered/*TP53*^WT^ (n=42), and *TP53-*altered/*KRAS*^WT^ (n=32) cohorts. To validate these findings, we queried: (1) The Cancer Genome Atlas (TCGA) PAAD non-silent somatic mutation data (v0.2.8) with survival data from Xena [14]; and (2) International Cancer Genome Consortium (ICGC) PACA-CA data set with survival information (https://dcc.icgc.org/).

### Transcriptomic analysis in TCGA and CCLE datasets

Overall transcriptome similarity of *KRAS*-*TP53* co-altered, *KRAS*-altered/*TP53*^WT^, and *TP53*-altered/*KRAS*^WT^ samples in the TCGA-PAAD dataset was examined using principal component analysis (PCA) based on log2-scaled RSEM values. For subsequent transcriptomic analysis, *KRAS-TP53* co-altered and *KRAS*-altered/*TP53*^WT^ TCGA-PAAD samples were utilized. Differential expression (DE) analysis of log2-transformed RSEM-normalized data matrices was performed using DESeq2 [15] for primary tumor samples; the “stat” column from DE results was used as input for Gene Set Enrichment Analysis (GSEA) using R package “fgsea”, and analyzed with Metascape knowledgebase (https://metascape.org) [16] or MSigDB:C2CP v6.2 [17] (“canonical pathways”) and Pan-Immune signature sets [18]. Complete GSEA results are included in **Table S2**.

TCGA-PAAD samples were stratified into consensus molecular [18] and consensus immune C1-C6 [18] subtypes, using TCGAbiolinks [19]. For canonical and Pan-Immune pathways common to C1+C2vs.C3 and *KRAS-TP53* co-altered vs. *KRAS*-altered, pathway overlap map was analyzed using EnrichmentMap [20] and visualized in Cytoscape [21]. Overlapping pathways common to DE genes in *KRAS-TP53* co-altered TCGA-PAAD samples and 9 stemness-related gene sets (**Table S3**) were visualized with Circos plot, generated using R-package “circlize” [22]. To identify predicted master regulators associated with innate immune subnetworks differentially expressed in TCGA-PAAD *KRAS-TP53* co-altered vs. *KRAS* -altered/*TP53*^WT^ genes (FDR-adjusted P<0.05), upstream regulator analysis were performed using Ingenuity Pathway Analysis (IPA; Qiagen Inc.) [23].

For analysis in human PDAC cell lines, genomic/transcriptomic sequencing data were retrieved from Cancer Cell Line Encyclopedia (CCLE; https://depmap.org/portal/ccle), and stratified into *KRAS-TP53* co-altered (n=32) and *KRAS*-altered/*TP53*^WT^ (n=8) (**Table S4**). Stratification into squamous or non-squamous (composite of progenitor, ADEX, and immunogenic) molecular subtypes was performed using TCGAbiolinks [19].

### Immune deconvolution analysis

Relative abundances of 24 immune cells (18 T-cell subtypes and 6 others—B-cell, NK-cell, monocyte, macrophage, neutrophil, and DC) using Immune Cell Abundance Identifier (ImmuCellAI [24]) were estimated in TCGA-PAAD samples, stratified by *KRAS/TP53* status. Curated immune-focused GSEA was also used to estimate immune populations, inflammation-related pathways, and allied immunologic pathways in indicated analyses (**Table S5**). Normalization Enrichment Score (NES) and FDR of each gene set was calculated using the “clusterProfiler”(v3.12.0) package [25].

### Cancer stemness index (mRNASi) annotation

Transcriptomes from TCGA-PAAD and CCLE samples, stratified by *KRAS-TP53* genomic status, were assigned mRNASi [26], by using the function TCGAanalyze_Stemness from the package TCGAbiolinks [19] following our previously described workflow. For details, refer to **Supplementary Methods**.

### scRNAseq datasets and analysis

scRNAseq datasets for *Kras*-*Trp53* co-altered “Late KPC” (GEO:GSM3577885) and *Kras*-altered/*Trp53*^WT^ “Late KIC” (GEO:GSM3577884) [27] were retrieved from NCBI. Panc02 scRNAseq data [28] were retrieved with permission (Dr. Huang) from Sequence Read Archive NCBI repository (#SRX7873760). Seurat v3.2 was used for cluster identification, and output visualized using tSNE/UMAP projections. For details, refer to **Supplementary Methods**.

Using R-package Monocle v2.8.0 [29], a differentiation hierarchy within the Panc02 granulocytic compartment was reconstructed. Using unbiased clustering, the top-20 unique genes per cluster were used to order cells along a pseudo-temporal trajectory. The cell sub-clusters obtained in Monocle were exported and compared to the single-cell lineage states predicted by RNA velocity using velocyto package [30]. Normalization and clustering were performed using PAGODA2 [31].

### In-vitro and in-vivo experiments

KC-PDA4313 (*Kras*-altered/*Trp53*^WT^; courtesy Andrew Lowy/UCSD), KPC-6694c2 or KPC^EV^ (*Kras-Trp53* co-altered), and KPC cells in which *Cxcl1* was genetically silenced using CRISPR/Cas9 gene editing technology KPC-*Cxc1*^KO^ (latter two courtesy Ben Stanger/UPenn [32]) and were used for *in-vitro* and *in-vivo* studies. Mutant *TP53*^R175H^ or *TP53*^WT^ cDNA constructs were transiently overexpressed in human HPNE-*KRAS*^G12D^ cells (ATCC).

All animal experiments were performed in accordance with the NIH animal use guideline and protocol approved by the Institutional Animal Care and Use Committee (IACUC) at UMiami. Tumors derived from KC and KPC GEMMs were used as indicated. For details of all *in-vitro* and *in-vivo* procedures, refer to **Supplementary Methods**.

### Generation of KRAS-TP53 IRP signature

Following concatenation of 20 transcripts differentially expressed in *KRAS-TP53* co-altered PDAC into a *KRAS-TP53* IRP signature, we assigned each sample from COMPASS, ICGC, and TCGA datasets an IRP score—i.e., number of genes (of 20) per sample whose expression was >median expression of the individual gene in the respective dataset. Subjects were dichotomized into a “high” (score>10) and “low” IRP (score<10; score=10 excluded), and compared via log-rank test in Kaplan-Meier survival analysis.

### COMPASS trial dataset and chemoresistance signatures

RNA sequencing data, basal-like/classical subtype classification, chemotherapy response, and survival information for patients accrued in the COMPASS trial (n=195) [33, 34] were provided by Ontario Institute for Cancer Research, and retrieved from EGAD00001006081. Chemotherapy response was available for 160 patients and dichotomized into progressive disease and partial response/stable disease.

The marker gene sets for chemotherapy resistance signatures were obtained from the literature [35-38], and single-sample GSEA [39] was used to quantify the NES of each chemoresistance signature, dichotomized into high and low *KRAS-TP53* IRP expression.

### Statistical analysis

For *in-vitro*/*in-vivo* experiments and clinical variables, two-tailed *t*-tests were used to compare continuous variables, Fisher’s exact test for categorical variables, and one-way ANOVA with Bonferroni post-hoc comparison for multi-level comparison. For DE gene/pathway analyses, features with FDR-adjusted P-values≤0.05 were considered statistically significant. OS was examined from date of diagnosis to date of death or last available follow-up. Kaplan-Meier survival curves were generated and compared using the log-rank test. Multivariable Cox regression was performed in the UMiami dataset using a forward stepwise methodology. Statistical analyses were performed using R v3.4.1 (https://cran.r-project.org/) and GraphPad Prism v8 (La Jolla, CA).

## RESULTS

### KRAS-TP53 co-alteration is associated with worse survival and immune exclusion in PDAC patients

We stratified samples in UMiami, TCGA, and ICGC datasets into genomic subgroups: *KRAS-TP53* co-altered, *KRAS*-altered/*TP53*^WT^, *TP53*-altered/*KRAS*^WT^, or *KRAS-TP53*-wildtype. First, a pan-cancer TCGA analysis revealed the highest prevalence of *KRAS-TP53* co-occurrent alterations in pancreatic adenocarcinoma (PAAD; 54%), followed by rectal (READ; 31%) and colon adenocarcinoma (COAD; 26%), and uterine carcinosarcoma (UCS; 12%) (**Fig 1A**).

**Figure 1:**
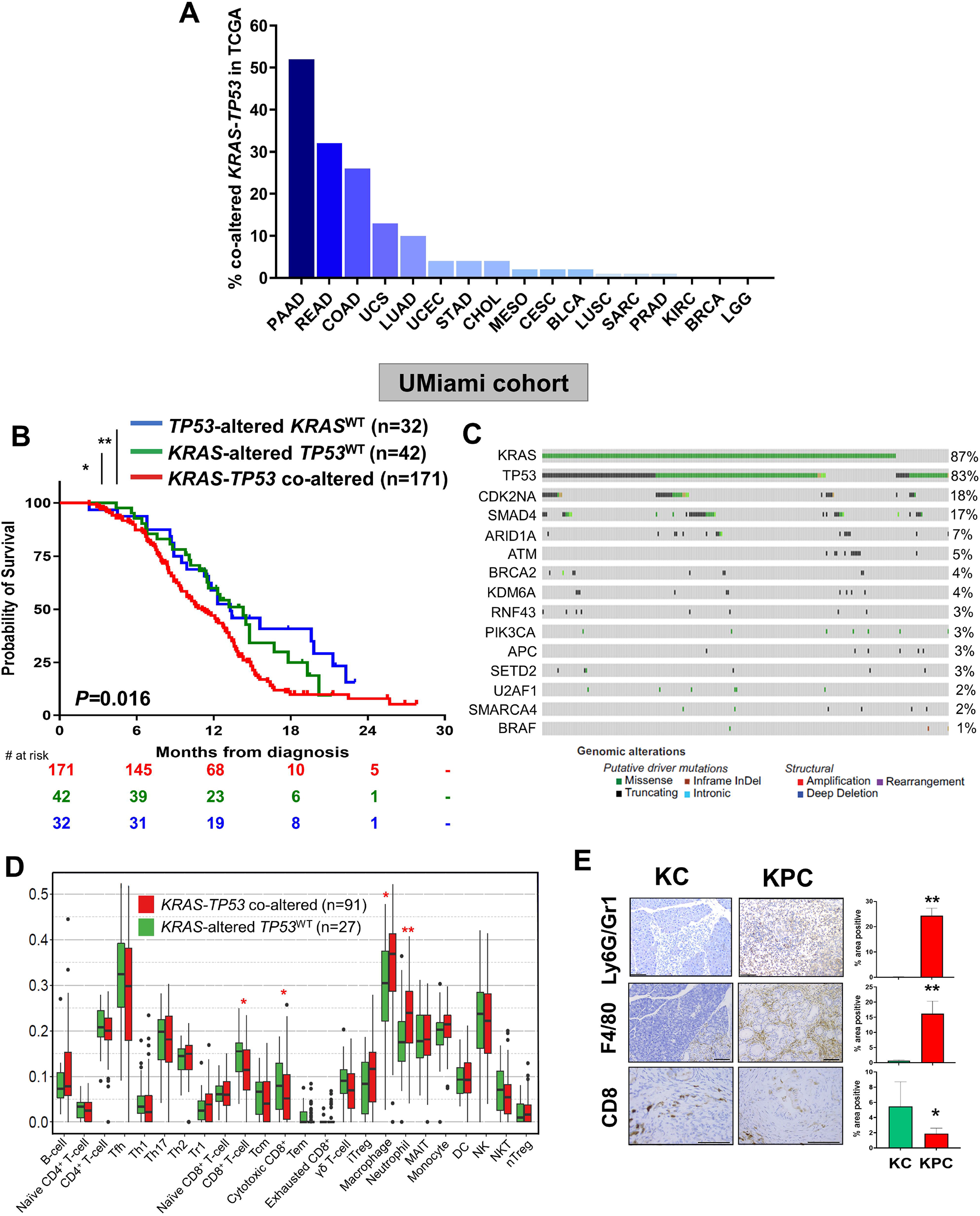
*KRAS-TP53* genomic co-alteration is associated with worse survival and immune exclusion in patients with pancreatic cancer. **(A)** Histogram showing proportion of pan-cancer samples from The Cancer Genome Atlas (TCGA) with co-occurrent alterations in *KRAS* and *TP53* [PAAD: Pancreatic adenocarcinoma, READ: Rectum adenocarcinoma, COAD: Colon adenocarcinoma, UCS: Uterine Carcinosarcoma, LUAD: Lung adenocarcinoma, UCEC: Uterine Corpus Endometrial Carcinoma, STAD: Stomach adenocarcinoma, CHOL: Cholangiocarcinoma, MESO: Mesothelioma, CESC: Cervical squamous cell carcinoma and endocervical adenocarcinoma, BLCA: Bladder Urothelial Carcinoma, LUSC: Lung squamous cell carcinoma, SARC: Sarcoma, PRAD: Prostate adenocarcinoma, KIRC: Kidney renal clear cell carcinoma, BRCA: Breast cancer, LGG: Brain lower grade glioma]. Only tumors with at least one *KRAS*-*TP53* co-altered sample (n=17) are included in the histogram; **(B)** Kaplan-Meier curves depicting overall survival of advanced PDAC patients (n=245) receiving treatment at the University of Miami, stratified by the presence of *KRAS-TP53* co-altered (n=171), *KRAS* - altered/*TP53*^WT^ (n=42), and *TP53*-altered/*KRAS*^WT^ (n=32) status. Adjoining table details number of patients at risk at designated time points; **(C)** Oncoprint representation of targeted next-generation genomic sequencing of advanced PDAC patients treated at the University of Miami (n=245), highlighting the 15 most frequent putative driver alterations. Types of gene alteration grouped by putative driver mutations or structural alterations are shown in the adjoining color legend. Somatic alteration frequencies in recurrently altered genes corresponding to specific genes are shown alongside; **(D)** Immune deconvolution using Immune Cell Abundance Identifier estimating relative abundance of 24 immune subpopulations in in TCGA PDAC samples (n=118), stratified by *KRAS-TP53* co-altered (red; n=91) and *KRAS*-altered/*TP53*^WT^ (green; n=27) tumors; **(E)** Representative photomicrographs of pancreatic tissue sections with adjoining quantification comparing the relative abundance of tumor-infiltrating Ly6G/Gr1^+^ myeloid cells; F4/80^+^ macrophages; CD8^+^ T-cells by immunohistochemical staining in murine pancreatic tumors (n=3 each) from *LSL-K-ras*^*G12D/+*^*;Pdx-1*^*Cre*^ (KC; 12-14 months of age) and *LSL-K-ras*^*G12D/+*^; *Trp53*^*R172H/+*^; *Pdx-1*^*Cre*^ (KPC) genetically engineered mice (5-6 months of age). \**p* <0.05; \*\**p* <0.01.

In 245 patients with metastatic PDAC receiving treatment at UMiami and undergoing targeted genomic sequencing, median age was 68 (range:32-94) years, 53% were male, 93% presented with synchronous metastatic disease, 14% had more than one distinct site of metastasis at diagnosis, and 75% had multi-agent chemotherapy exposure prior to sequencing (**Table S1**). In this population of exclusively metastatic PDAC patients, at a median follow-up of 11.1 months (range 2.3-27.8), median, 1- and 2-year overall survival (OS) was 12.2 months, 44%, and 2% respectively. Importantly, patients with *KRAS-TP53* co-alteration (n=171) had significantly worse median OS compared with patients with *KRAS*-altered/*TP53*^WT^ (n=42) or *TP53*-altered/*KRAS*^WT^ (n=32) tumors (median 11.1 vs. 14.2 vs. 13.3 months, respectively; P=0.016) (**Fig 1B**). Multivariable Cox regression confirmed the independent prognostic impact of *KRAS-TP53* co-alteration (HR 2.1, 95%CI 1.2-5.6, P=0.002) and older age (HR 1.9, 95%CI 1.2-4.7, P<0.001) while controlling for gender, stage at diagnosis, and chemotherapy regimen (**Table S1**). Beyond *KRAS* or *TP53* gene alterations, the most common putative driver alterations were observed in *CDK2NA* (18%), *SMAD4* (17%), and *ARID1A* (7%) (**Fig 1C**).

We validated the prognostic impact of *KRAS-TP53* genomic co-alteration in two independent datasets. In the ICGC dataset comprising patients with both localized and metastatic PDAC (n=230), patients with *KRAS-TP53* co-alteration (n=173) had significantly worse median OS compared with patients with *KRAS*-altered/*TP53*^WT^ (n=46) or *TP53*-altered/*KRAS*^WT^ (n=11) tumors (median 16 vs. 25 vs. 27 months, respectively; P=0.005) (**Fig S1A**). In the TCGA dataset (n=135), patients with *KRAS-TP53* co-altered PDAC (n=91) demonstrated worse survival compared with patients with *KRAS*-altered/*TP53*^WT^ (n=27) or *TP53*-altered/*KRAS*^WT^ PDAC (n=17) (median 16 vs. 22 vs. 31 months, respectively; P=0.045) (**Fig S1B**).

We have previously described the relevance of *KRAS-TP53* genomic co-alteration as a model to investigate mechanisms of immune regulation in non-PDAC GI malignancies [7]. Comparing the abundance of computationally-inferred populations via immune deconvolution [24] in TCGA-PAAD samples revealed significant increases in tumor-associated neutrophils and macrophages, with concomitant reduction in global and cytotoxic CD8^+^ T-cell populations in *KRAS-TP53* co-altered compared with *KRAS*-altered/*TP53*^WT^ tumors (**Fig 1D**). To confirm these findings, we examined volume-matched tumor sections from *Kras*-altered/*Trp53*^WT^ *LSL-K-ras*^*G12D/+*^*;Pdx-1*^*Cre/+*^ (KC; 12-14 months of age) and *Kras-Trp53* co-altered KPC (5-6 months of age) PDAC GEMMs *in-vivo*. Compared with KC lesions (i.e., PanIN3/early invasive carcinoma), KPC tumors (i.e., early invasive carcinoma) were more abundantly infiltrated by Ly6G^+^ tumor-infiltrating neutrophils—by nomenclature synonymous with and phenotypically similar to Gr1^+^Ly6G^+^ granulocytic myeloid-derived suppressor cells (gMDSCs) [40]—and F4/80^+^ macrophages, and relatively deficient in CD8^+^ T-cells (**Fig 1E**). Taken together, these data indicate that *KRAS-TP53* co-altered PDAC is not only associated with worse survival, but is also innate immune-enriched and CD8^+^ T-cell-excluded compared with *KRAS*-altered PDAC.

### The KRAS-TP53 co-altered transcriptome intersects with squamous transdifferentiation and immune exclusion in PDAC

Our subsequent investigations focused on comparing the differentially expressed transcriptomes in *KRAS-TP53* co-altered with *KRAS*-altered/*TP53*^WT^ human PDAC for several reasons: (a) Mutated *KRAS* is the incipient genomic alteration in >90% of PDAC, representing a foundational molecular “baseline” exploited by other oncogenic driver alterations (e.g., *TP53*) which exert co-operative tumor-permissive signaling [8, 10, 41-43]; (b) *TP53*-altered/*KRAS*^WT^ PDAC comprises <5% of human disease [44]; (c) only 11 *TP53*-altered/*KRAS*^WT^ tumors with clinically annotated transcriptomic data in the TCGA-PAAD dataset precluded a powered three-way comparison; and (d) PCA analysis revealed *non-overlapping* transcriptomes between TCGA-PAAD *KRAS*-altered/*TP53*^WT^ and *TP53*-altered/*KRAS*^WT^ tumors—both distinct from *KRAS-TP53* co-altered transcriptomes—thereby precluding merging of these two datasets (**Fig S2A**).

Pathway analysis using MSigDB:C2CP or Metascape knowledgebase in DEGs from *KRAS-TP53* co-altered (n=91) compared with *KRAS*-altered/*TP53*^WT^ (n=27) TCGA-PAAD samples revealed that pathways related to p63/p73 signaling and ΔNP63 transcriptional targets were strongly enriched in *KRAS-TP53* co-altered PDAC (P-adj<0.001). In addition, pathways related to myeloid/innate immune activation, neutrophil degranulation/trafficking, cancer stemness and stem cell proliferation, and regulation of DNA damage response were among the top 20-enriched upregulated pathways (**Fig 2A, Fig S2B**). When constructing a protein-protein interactome (PPI) of the *KRAS-TP53* co-altered transcriptome using the STRING network, *TP63* was a significantly connected node to both KRAS and TP53 in this interactome (enrichment P<1.0e-16, suggesting significant biological connectivity; **Fig S2C**).

**Figure 2:**
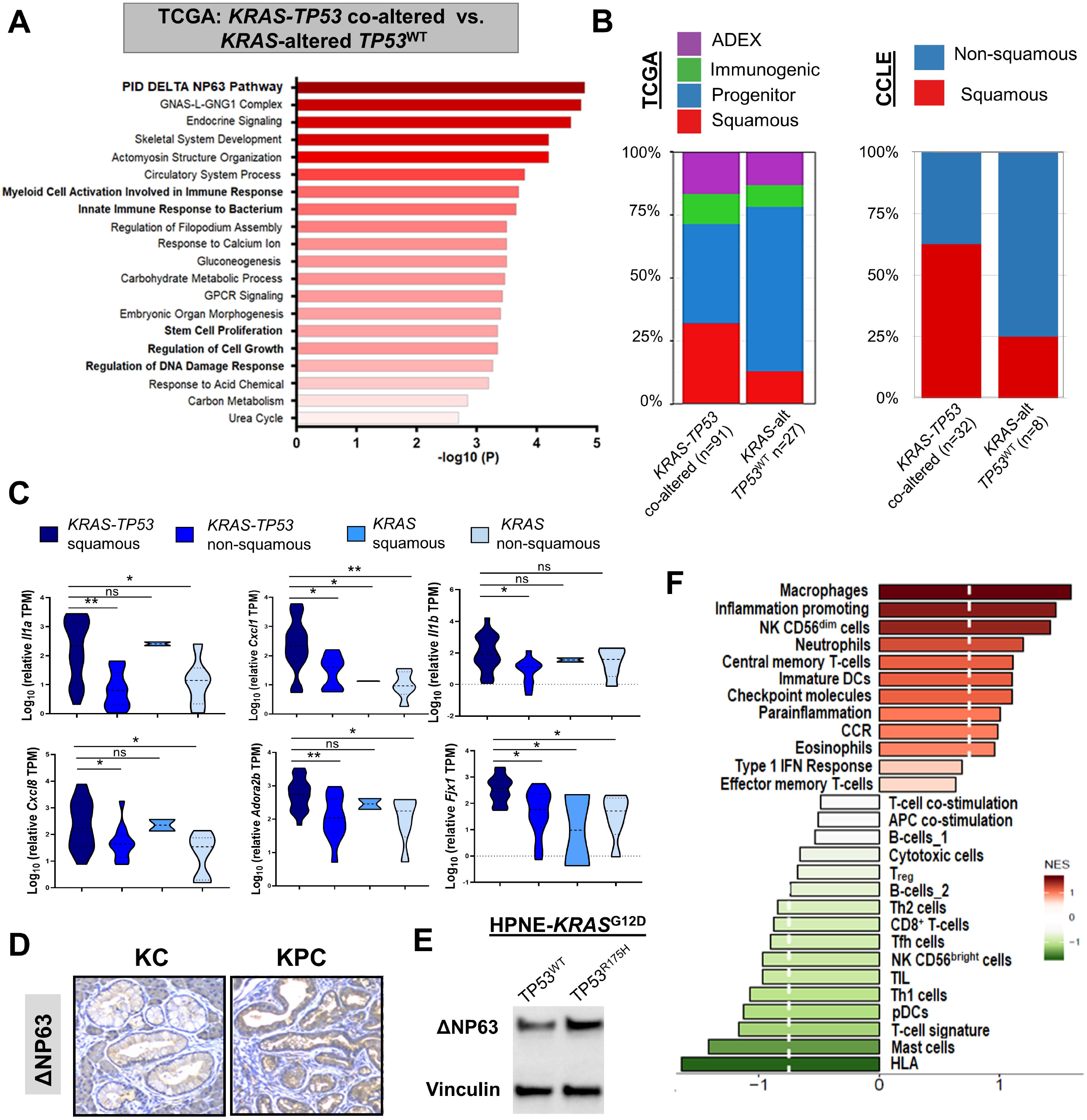
The *KRAS-TP53* co-altered transcriptome intersects with *TP63*-defined squamous transdifferentiation and immune exclusion in PDAC. **(A)** Metascape pathway enrichment analysis depicting canonical signaling pathways (C2:CP) from MSigDB compendium differentially upregulated in *KRAS-TP53* co-altered compared with *KRAS*-altered/*TP53*^WT^ TCGA PDAC samples. Bolded pathways highlight those related to ΔNP63 signaling, innate immunoregulation, and cancer stemness; **(B)** Stacked histograms representing the proportions of samples classified into (*left)* consensus molecular subtype (Squamous, Progenitor, Aberrantly Differentiated Endocrine Exocrine [ADEX], Immunogenic) across *KRAS-TP53* co-altered (n=91), and *KRAS* -altered *TP53*^WT^ (n=27) cohorts in TCGA dataset, and (*right*) proportions of Squamous and Non-Squamous subtypes, annotated using TCGABiolinks algorithm, in *KRAS-TP53* co-altered (n=32) and *KRAS*-altered *TP53*^WT^ (n=8) human PDAC cell lines in Cancer Cell Line Encyclopedia (CCLE); **(C)** Normalized expression (transcripts per million [TPM]) of six transcripts (*Il1a, Cxcl1, Il1b, Cxcl8, Adora2b, Fjk1)* implicated in squamous trans-differentiation in PDAC, stratified by *KRAS-TP53* co-alteration and squamous/non-squamous molecular subtype status; **(D)** Representative section depicting ΔNP63 expression by immunohistochemical staining in KPC (age 5-6 months) and KC (age 12-14 months) GEMM tumors; **(E)** ΔNP63 protein levels depicted by western blot in presence or absence of *TP53*^*R175H*^ mutant plasmid transfection in an immortalized epithelial human pancreatic-expressing cell line harboring *KRAS*^*G12D*^ mutation (HPNE-K-ras^G12D^); **(F)** Normalized enrichment scores of curated gene sets encompassing immunologic and inflammatory pathways evaluated using Gene Set Enrichment Analysis (GSEA) for differential expression in squamous versus non-squamous samples within *KRAS-TP53* co-altered TCGA-PAAD tumors.

Building on recent evidence implicating *TP63* as a master regulator of squamous trans-differentiation in PDAC [45], we observed enrichment of the squamous molecular subtype [46] in *KRAS-TP53* co-altered tumors compared with *KRAS*-altered/*TP53*^WT^ tumors in the TCGA-PAAD dataset (**Fig 2B**). Moreover, assigning consensus molecular subtypes to CCLE-derived human PDAC cell lines revealed significant enrichment of the squamous (vs. non-squamous) phenotype in *KRAS-TP53* co-altered (n=32) compared with *KRAS*-altered/*TP53*^WT^ (n=8) tumor-cell transcriptomes (χ^2^-P=0.03) (**Fig 2B, Table S4**).

Using TCGABiolinks-nominated subtype classification, stratification of CCLE PDAC tumor-cell lines into *KRAS-TP53* squamous (n=20), *KRAS-TP53* non-squamous (n=12), *KRAS*-altered/*TP53*^WT^ squamous (n=2), and *KRAS*-altered/*TP53*^WT^ non-squamous (n=6) revealed significant overexpression of squamous trans-differentiation-defining transcripts in PDAC—*Il1a, Cxcl1, Il1b, Cxcl8, Adora2b, Fjx1[45]*—in the *KRAS-TP53* squamous PDAC transcriptome (**Fig 2C**). These transcriptomic changes correlated with the increased secretion of CXCL1 and IL-1α from *KRAS-TP53* co-altered human and murine PDAC tumor-cell lines *in-vitro*, compared with *KRAS*-altered/*TP53*^WT^ PDAC cells (**Fig S2D**). Furthermore, multiple transcripts from pathways defining squamous lineages in non-PDAC solid tumors [47] were also differentially expressed in the *KRAS-TP53* co-altered vs. *KRAS*-altered/*TP53*^WT^ TCGA-PAAD transcriptome (**Fig S2E, Table S2**). Corroborating these findings in high-fidelity PDAC GEMMs, TP63 ΔNP63 isoform expression was increased in the epithelial/acinar compartment of KPC compared to KC tumors *in-vivo* (**Fig 2D**).

To recapitulate *KRAS-TP53* co-alteration in a human pancreatic epithelial cell system, we transiently overexpressed mutant *TP53*^R175H^ or *TP53*^WT^ cDNA constructs in HPNE-*KRAS*^G12D^ cells and demonstrated increased ΔNP63 expression in dual-mutant *KRAS*^G12D^*-TP53*^R175H^, compared with *KRAS*^G12D^*-TP53*^WT^, cells *in-vitro* (**Fig 2E**). To investigate the clinical repercussions of ΔNP63 signaling in PDAC, a gene set encompassing transcriptional targets of ΔNP63 (PID_DELTA_NP63_PATHWAY) was significantly enriched in PDAC patients with chemoresistant disease (i.e., progressive disease by RECIST criteria) compared to patients with chemoresponsive disease (stable disease or partial response) in patients enrolled in the COMPASS trial [34] (P-adj=0.002; **Fig S3A**). Finally, TCGA-PAAD tumors overexpressing transcriptional targets of ΔNP63 demonstrated worse overall survival compared with tumors under-expressing ΔNP63 targets (HR 1.6; P=0.024) (**Fig S3B**). Collectively, these data suggest a novel and clinically relevant link between *KRAS-TP53* co-alteration and *TP63*-defined squamous trans-differentiation in PDAC.

Next, curated immune-based GSEA (**Table S5**) *within* the *KRAS-TP53* co-altered TCGA-PAAD subset revealed disproportionate enrichment of innate immune (macrophages, neutrophils) pro-inflammatory (inflammation-promoting, parainflammation, etc.), and immune checkpoint signatures, with downregulation of antigen-presenting/processing (pDCs, HLA) and immune effector (CD8^+^ T-cells, TIL, Th1, NK) signatures in squamous compared with non-squamous *KRAS-TP53* co-altered PDAC (**Fig 2F**). These data expose the intersection between *KRAS-TP53* co-alteration, squamous trans-differentiation, and immune exclusion in PDAC.

### KRAS-TP53 co-alteration encodes a transcriptional program that orchestrates myeloid cell migration to the PDAC TME

Further investigation into the transcriptional determinants associated with myeloid cell activation/trafficking in *KRAS-TP53* co-altered PDAC (**Fig 1&2**) revealed overexpression of several well-characterized genes (**Fig 3A**) and Pan-Immune [18] pathways (**Fig S4A**) implicated in innate immunoregulation, granulocyte trafficking/degranulation, macrophage recruitment, and inflammasome signaling/activation. Significantly overexpressed transcripts in *KRAS-TP53* co-altered—compared with *KRAS*-altered/*TP53*^WT^—PDAC overlapped with prognostically relevant immune gene signatures derived from human cancer tissue transcriptomes using a network-based deconvolution approach (*ImSig*) [48], and nominated macrophage-(*LY96, CD300A, FCGR2A, SCPEP1, CD163*), neutrophil-(*THBD, AQP9, NAMPT*), and inflammasome-related (*IL18, PANX1*) genes (**Fig 3B**).

**Figure 3:**
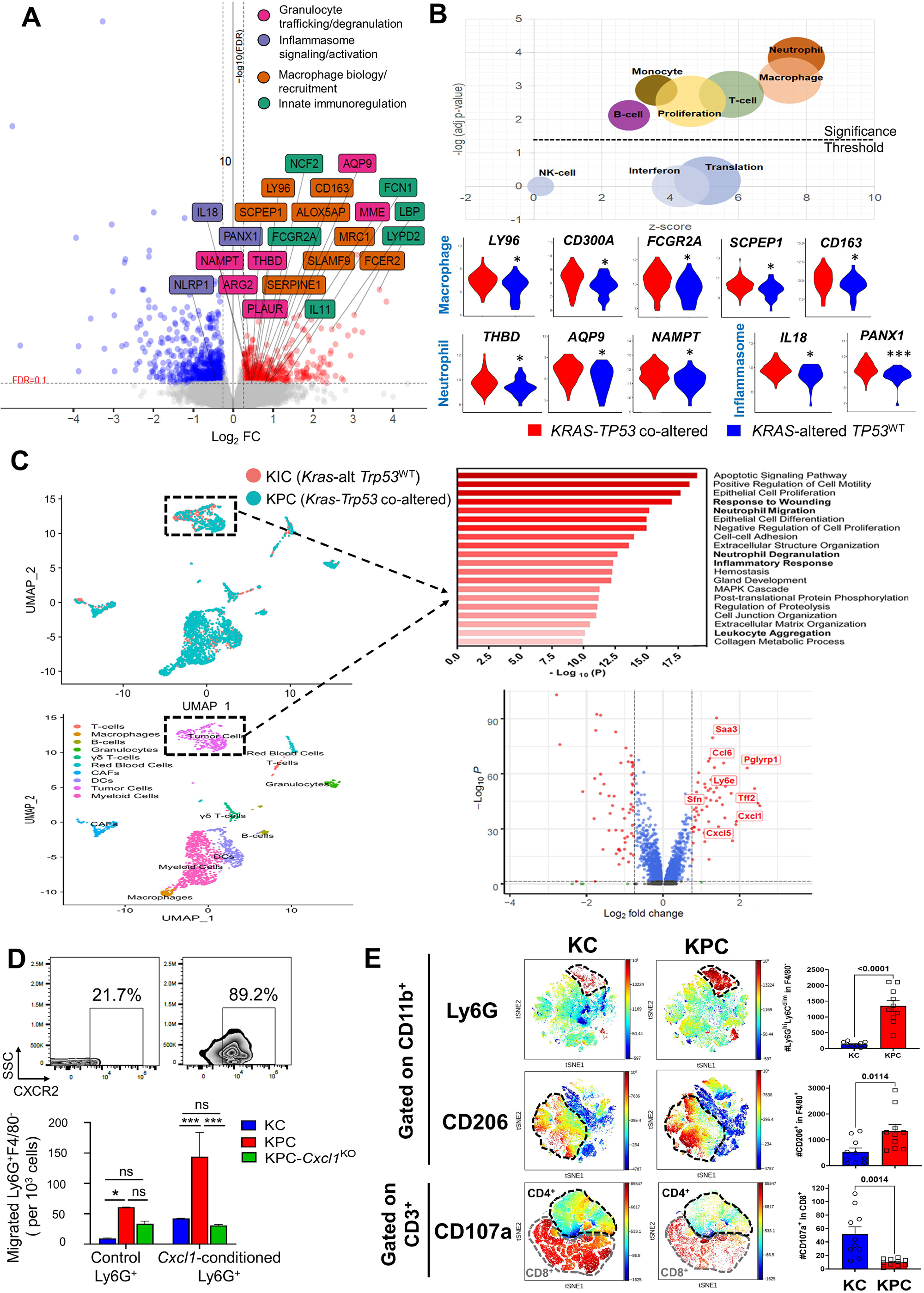
*KRAS-TP53* co-alteration encodes a transcriptional program that orchestrates myeloid cell migration and infiltration into the PDAC tumor microenvironment. **(A)** Volcano plot of innate immune pathway-related genes that are significantly overexpressed (*red)* or underexpressed (*blue*) in TCGA-PAAD *KRAS-TP53* co-altered compared to *KRAS*-altered *TP53*^WT^ PDAC, nominating genes implicated in innate immunoregulation, granulocyte trafficking/degranulation, macrophage recruitment, and inflammasome signaling/activation (*top right*); **(B)** (*top*) Bubble plot depicting an overlap between significantly overexpressed transcripts in *KRAS-TP53* co-altered PDAC and prognostically relevant immune gene signatures derived from human cancer tissue transcriptomes using a network-based deconvolution approach (*ImSig*). Gene sets are plotted on z-score, calculated as the difference between number of upregulated and downregulated genes over the square-root of the total number of genes belonging to the gene sets, against FDR-adjusted P-value with threshold of 0.05 (dashed line); (*bottom*) Several macrophage-, neutrophil-, and inflammasome-related genes nominated by *ImSig* are differentially overexpressed in *KRAS-TP53* co-altered (*red*) vs. *KRAS*-altered *TP53*^WT^ (*blue*) TCGA-PAAD samples, as depicted by violin plots; **(C)** Uniform Manifold Approximation and Projection (UMAP) of scRNAseq of combined KPC (*green*) and KIC (*orange*) datasets (*top left*), with annotated clustering map showing 9 distinct cellular constituents (*bottom left*). Within the tumor-cell compartment (indicated by dashed box), significantly upregulated genes in KPC vs. KIC datasets nominated pathways related to neutrophil migration and degranulation, leukocyte aggregation, and inflammatory response/wound healing (*top right*). Volcano plot depicts cancer cell-autonomous transcripts implicated in innate immune function/signaling significantly upregulated in KPC vs. KIC, particularly *Cxcl1, Cxcl5, Tff2, Sfn, Ly6e, Ccl6, Pglyrp1*, and *Saa3* (*bottom right*); **(D)** Pseudo-color flow cytometry plots showing proportion of CXCR2^+^ Ly6G^+^F4/80^-^ cells following *ex vivo* conditioning of C57/B6 murine bone marrow-derived cells with and without CXCL1—a strong neutrophil chemoattractant. Histogram visualizing Ly6G^+^F4/80^-^ (×10^3^) cells migrated through the transwell insert (6h timepoint) in presence of conditioned media from KC, KPC, or KPC tumor cells in which CXCL1 is genetically silenced using CRISPR/Cas9 editing technology (KPC-*Cxcl1*^KO^); **(E)** viSNE maps comparing Ly6G^+^F4/80^-^ gMDSC, and F4/80^+^CD206^+^ M2-like macrophage populations gated on CD11b^+^ cells, as well as CD107^+^ populations gated on CD3^+^ T-cells between volume-matched KPC vs. KC orthotopic tumors *in vivo* (n=10 mice/group). Dotted lines highlight the cell subset of interest in each viSNE plot. Adjoining histograms show absolute number of relevant cell populations per 1×10^4^ live cells; \**p* <0.05; \*\*\**p <*0.001.

Next, to explore cancer cell-*autonomous* transcriptional programs that orchestrate innate immune trafficking and function, we compared single-cell transcriptomes in the tumor-cell clusters between *Kras-Trp53* co-altered KPC and *Kras*-altered/*Trp53*^WT^ KIC datasets [27]. This analysis confirmed significant enrichment of pathways related to neutrophil migration and degranulation, leukocyte aggregation, and inflammatory/wound healing response, and nominated a *Kras-Trp53* co-altered tumor-cell-*intrinsic* transcriptional program—comprising *Cxcl1, Saa3, Cxcl5, Pglyrp1, Ly6e, Tff2, Sfn*, and *Ccl6*—which promotes a neutrophil/myeloid-infiltrated immune contexture (**Fig 3C, Fig S4B**).

Next, to functionally assess the comparative effects of *Kras-Trp53* co-altered vs. *Kras*-altered/*Trp53*^WT^ PDAC on granulocyte migration *ex-vivo*, we performed migration assays in which murine granulocytic MDSCs (gMDSC) derived from spleens of tumor-bearing KPC mice—with or without exogenous conditioning with CXCL1—were co-cultured with either *Kras-Trp53* co-altered (KPC-6694c2) or *Kras*-altered/*Trp53*^WT^ (KC-PDA4313) cells in transwell systems. While the KPC secretome enhanced the transwell migration of non-CXCL1 conditioned gMDSCs compared to the KC secretome at baseline, CXCL1 conditioning of gMDSCs not only augmented the expression of its cognate receptor CXCR2 >4-fold, but also further accentuated the disparate effects of KPC-induced transwell migration of CXCR2^hi^ gMDSCs *ex-vivo*. These gMDSC migratory effects were also dependent on cancer cell-intrinsic CXCL1, since CRISPR/Cas9 genetic silencing of *Cxcl1* in KPC tumor cells (KPC-*Cxcl1*^KO^ [32]) significantly abrogated CXCR2^hi^ gMDSC trafficking compared with KPC tumor secretomes (**Fig 3D**).

Next, multiparameter flow cytometric immunophenotyping of orthotopic KC-PDA4313 and KPC-6694c2 tumors *in vivo* (n=10 mice/group) revealed disproportionately increased infiltration of CD11b^+^ myeloid cells, F4/80^-^Ly6G^hi^Ly6C^dim^ gMDSCs, F4/80^+^ macrophages, and specifically F4/80^+^CD206^+^ M2-like (but not F4/80^+^CD86^+^ M1-like) macrophages into KPC compared with KC TMEs (all P<0.05; **Fig 3E, Fig S4C**). This myeloid-enriched TME was also associated with significantly decreased CD8^+^CD107a^+^ degranulating cytotoxic T-cells in KPC vs. KC tumors *in-vivo* (P<0.001; **Fig 3E**). Taken together, these data reveal novel cancer cell-autonomous transcriptional programs encoded by *Kras-Trp53* co-alteration that functionally regulate intratumoral neutrophil/myeloid migration and trafficking into the PDAC TME.

### Granulocyte-derived inflammasome activation and TNF signaling are putative paracrine mediators of innate immunoregulatory transcriptional networks in KRAS-TP53 co-altered PDAC

We next explored putative non-cancer cell-autonomous mechanisms that mediate innate immune regulation in *KRAS-TP53* co-altered PDAC. To build on our observation that inflammasome signaling was differentially upregulated in *KRAS-TP53* co-altered PDAC (**Fig 3A&B**), interrogation of the relative expression of the inflammasome transcriptional machinery in single-cell transcriptomes across key cellular clusters from *Kras-Trp53* co-altered vs. *Kras*-altered/*Trp53*^WT^ scRNAseq datasets revealed significant overexpression of *Nlrp3* and *Il18* in tumor-cell, cancer-associated fibroblast (CAF), and granulocyte/gMDSC, but not monocytic/macrophage, clusters in *Kras-Trp53* co-altered tumors (**Fig 4A**). To compare *functional* inflammasome activation in these cellular subsets, flow cytometry-sorted F4/80^-^Ly6G^+^ granulocyte/gMDSCs, monocytic, PDPN^+^ CAFs, and EpCAM^+^ tumor-cells from KPC and KC tumor-bearing mice subjected to caspase-1 luminescence assays revealed significantly increased inflammasome activation exclusively in the granulocyte/gMDSC, but not tumor-cell, CAF, or monocytic compartments (**Fig 4B**). These results were corroborated using flow cytometry showing disproportionately higher IL-1β expression—the terminal product of inflammasome activation—in KPC vs. KC tumor-infiltrating gMDSC (P<0.001), but not in EpCAM^+^ tumor cells, PDPN^+^ CAFs, or Ly6C^+^ monocytic cells (P=ns; **Fig S5A**).

**Figure 4:**
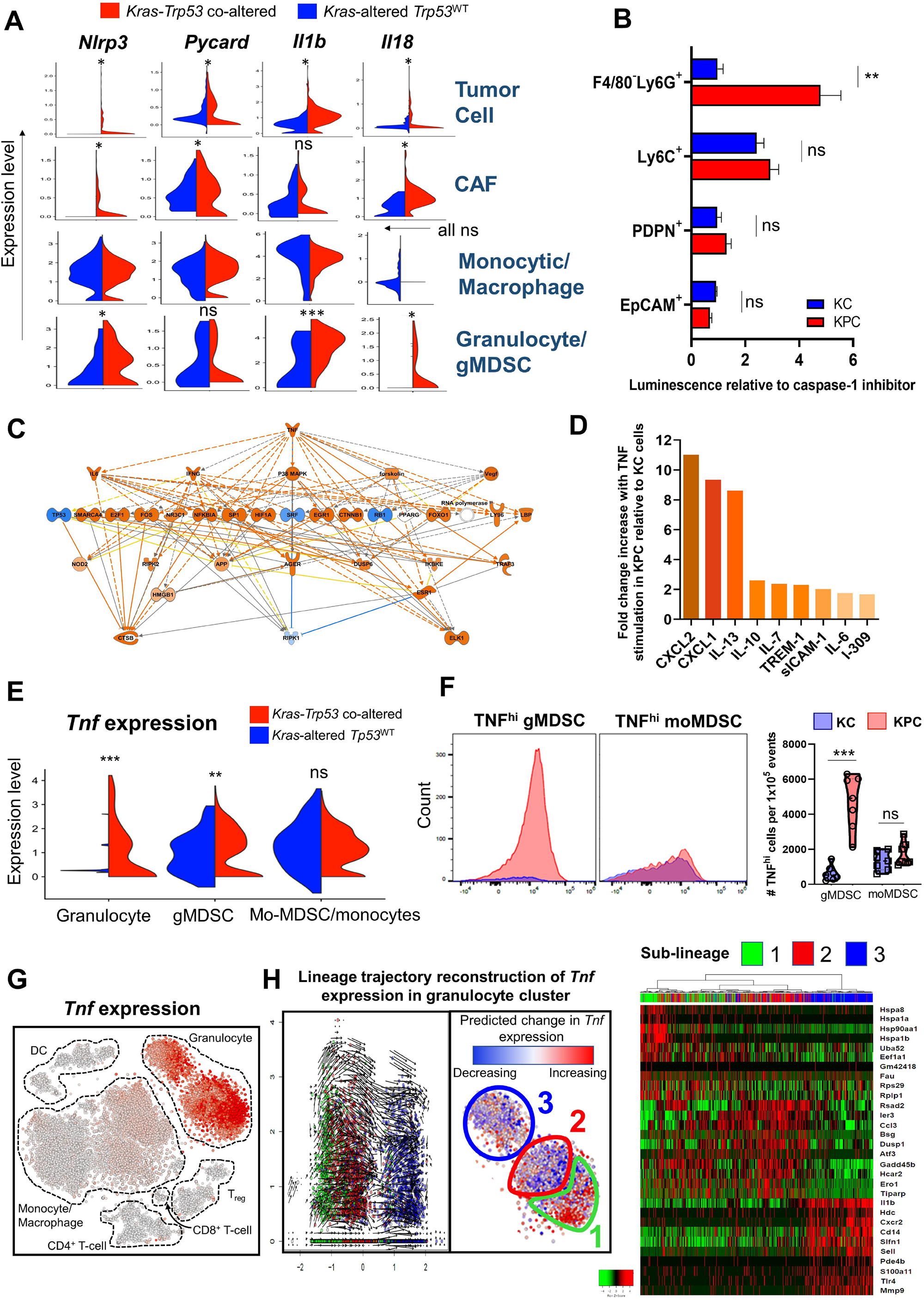
Granulocyte-derived inflammasome and TNF activation are putative mediators of immunoregulation in *KRAS-TP53* co-altered PDAC. **(A)** Violin plots showing the differential expression of inflammasome machinery-related genes (*Nlrp3, Pycard, Il1b, Il18*) in KPC compared with KIC scRNAseq datasets in tumor-cell, cancer-associated fibroblast (CAF), monocytic/macrophage, and granulocytic myeloid-derived suppressor cells (gMDSC)/granulocyte clusters; **(B)** Caspase-1 functional inflammasome activity assay in flow cytometry-sorted F4/80^-^Ly6G^+^ (gMDSC), Ly6C^+^ (monocyte), PDPN^+^ (CAF), and EpCAM^+^ (tumor-cell) populations derived from KPC and KC orthotopic tumors (n=4-5 mice/group) are compared in histograms. Plotted values for each group represent luminescence relative to caspase-1 inhibitor treatment control; **(C)** Predicted upstream regulators of differentially expressed genes that comprise the innate immune subnetwork in TCGA-PAAD *KRAS-TP53* co-altered vs. *KRAS* -altered *TP53*^WT^ PDAC samples, by Ingenuity Pathway Analysis (IPA); **(D)** Histogram visualizing relative fold increase in secretion of cytokines associated with innate immune trafficking/function after 48 hours of exogenous TNF conditioning (50ng/ml) in KPC-6694c2 tumor cells compared with KC-PDA4313 cells; **(E)** Violin plots representing differential *Tnf* gene expression among granulocytic, gMDSC, and monocytic MDSC/monocyte sub-clusters in KPC (*red*) versus KIC (*blue*) scRNAseq datasets; **(F)** Representative overlayed histogram plots highlighting relative counts of high TNF-expressing gMDSC (Ly6G^+^Ly6C^lo^) and moMDSC (Ly6G^-^Ly6C^hi^) infiltrating KPC (*left)* vs. KC (*right*) orthotopic tumors *in vivo*, and adjoining violin plots showing the quantification across mice in each group (n=10 mice/group); **(G)** tSNE feature plot of *Tnf* gene expression (*red*) in cellular sub-clusters of *KRAS-TP53* co-altered Panc02 tumor-infiltrating CD45^+^ cells by scRNAseq, demonstrating highest *Tnf* expression in granulocyte cluster; **(H)** Within the granulocyte cluster, lineage trajectory reconstruction by RNA velocity algorithm depicts differences in *Tnf* splicing among three distinct granulocytic sub-lineages. Cells were projected to the line y=-x centered to medians of the extracted granulocyte clusters and ordered from left to right (unspliced➔spliced transcripts) along the X-axis, and plotted against expression levels of *Tnf* (Y-axis; *left*). The predicted tendency of *Tnf* transcription towards a future state (increasing=red; decreasing=blue) by RNA velocity analysis is depicted in the granulocytic cluster. Monocle-enabled pseudotemporal sub-lineage prediction was superimposed on this RNA velocity analysis, as illustrated by outlines and denoted by sub-lineages 1-3 (*middle*). Heatmap depicts the top 20-expressed genes per granulocytic sub-lineage as predicted by Monocle pseuodotemporal ordering (*right)*. ns: non-significant; \**p* < 0.05; \*\**p* < 0.01; \*\*\**p <*0.001.

In exploring upstream regulators via Ingenuity Pathway Analysis (IPA) that control innate immune transcriptional subnetworks differentially expressed in *KRAS-TP53* co-altered TCGA-PAAD samples, *TNF* emerged as the predicted candidate (**Fig 4C**). To simulate this functionally *in-vitro*, exogenous TNF conditioning of KPC-6694c2 and KC-PDA4313 tumor cells resulted in substantial relative overproduction of granulocytic and myeloid chemoattractant chemokines— including CXLC2, CXCL1, IL-13, TREM-1, and IL-6—from KPC tumor-cells (**Fig 4D**). Next, examination of myeloid single-cell transcriptomes from *Kras-Trp53* co-altered vs. *Kras*-altered/*Trp53*^WT^ datasets (**Fig S5B**) revealed significant overexpression of *Tnf* in granulocytic and gMDSC, but not monocytic/monocytic-MDSC (**Fig 4E**) cellular clusters. These results were validated in volume-matched KC and KPC orthotopic tumor-bearing mice using flow cytometry, revealing disproportionately higher intracellular TNF expression in gMDSCs—but not monocytic-MDSCs—derived from KPC compared with KC mice in both intratumoral (**Fig 4F**) and splenocytic (**Fig S5C**) compartments (both P<0.001).

In single-cell transcriptomes of CD45^+^ immune cells infiltrating *Kras-Trp53* co-altered Panc02 murine tumors [28], the granulocytic cluster—compared against monocytic/macrophage, dendritic cell, CD4^+^ and CD8^+^ T-cell, and regulatory CD4^+^ T-cell clusters—demonstrated the strongest expression of *Tnf* (**Fig 4G**). Next, in order to explore novel molecular markers associated with *Tnf* gene expression in this Panc02-associated granulocyte cluster, we harmonized two established pipelines (i.e., RNA velocity [30] and Monocle [29]) to reconstruct *Tnf* transcriptional kinetics across distinct cellular developmental trajectories. In RNA velocity analyses, the fraction of spliced *Tnf* transcripts (i.e., only exons) represents the current state of the cell, while the fraction of unspliced *Tnf* transcripts defines an RNA velocity vector predicting the future state of the cell (**Fig 4H**; *left*). Superimposing Monocle-enabled pseudotemporal sub-lineage prediction on this RNA velocity analysis revealed granulocyte differentiation states with distinct transcriptional identities. The steady state of *Tnf* transcription was concentrated in a more phenotypically “mature” sub-lineage (sub-lineage 3), defined by expression of *Cxcr2, Il1b, Hdc*, and *Cd14*. Conversely, the predicted transcriptional tendency towards the future state of *Tnf* expression was largely confined to a phenotypically “immature” sub-lineage (sub-lineage 1), enriched for heat-shock protein family members *Hspa8, Hsp90aa1, Hspa1a*, and *Hspa1b* (**Fig 4H**; *right*). Taken together, these data identify inflammasome activation and *TNF* signaling as putative granulocyte-derived paracrine mediators of innate immune regulation in *KRAS-TP53* PDAC.

### T-cell exclusion in KRAS-TP53 co-altered PDAC is associated with distinct immunomodulatory and adaptive immune vulnerabilities

In parallel with the disproportionate innate immune trafficking and function in *Kras-Trp53* co-altered—compared with *Kras*-altered/*Trp53*^WT^—PDAC, we observed striking exclusion of CD4^+^, CD8^+^, and activated CD69^+^ in orthotopic KPC versus KC tumors *in-vivo* (**Fig S6A**). Exploration of the differentially expressed transcriptomes in *KRAS-TP53* co-altered vs. *KRAS* - altered/*TP53*^WT^ TCGA-PAAD datasets revealed a limited repertoire of transcripts associated with inhibitory adaptive immune function, such as *CD52* [binding to Siglec-10 receptor attenuates effector T-cell activation [49]], *EVI2A* [marker of inhibitory naïve CD4^+^ T-cells [50]], *MSN*, and *DSC3* (**Fig S6B**). Examining the differentially expressed genes in *KRAS-TP53* co-altered TCGA-PAAD in the context of a curated compendium of immunomodulatory genes [ImmunoModulator-78 [18]] revealed significant overexpression of *IL2RA* [51] and *MICB [52]*, and under-expression of *HLA-DQB2* and *TNFSF9* (i.e., co-stimulatory 4-1BB ligand) in *KRAS-TP53* co-altered PDAC (**Fig S6C**). Collectively, these data highlight unique molecular determinants in *KRAS-TP53* co-altered PDAC indicative of adaptive immune dysfunction and immune evasion.

### Immune subtyping reveals conflation of intratumor heterogeneity with augmented stemness properties in KRAS-TP53 co-altered PDAC

To gain further insight into the unique immunologic architecture of *KRAS-TP53* co-altered PDAC, we categorized TCGA-PAAD samples—stratified by *KRAS-TP53* alteration status—into six consensus immune subtypes (C1-C6), characterized by differences in extent of intratumoral heterogeneity (ITH), cell proliferation, neoantigen load, myeloid/lymphocyte signatures, Th1:Th2 ratio, expression of immunomodulatory genes, and prognosis [18]. *KRAS-TP53* co-altered PDAC tumor samples were highly enriched in C1 (wound healing) and C2 (IFN- γ dominant) subtypes, both associated with highest ITH and cellular proliferation, and low Th1:Th2 ratio. The incidence of the C3 (inflammatory) subtype—defined by lowest ITH and high Th1:Th2 ratio—was lowest in *KRAS-TP53* co-altered PDAC, and its prevalence increased progressively in *KRAS*-altered/*TP53*^WT^, *TP53*-altered/*KRAS*^WT^, and *KRAS-TP53*-wildtype tumors (**Fig 5A**). Of note, C1/C2 subtypes are associated with poor prognosis, while C3 subtype confers the best prognosis in pan-cancer analysis [18]. Mapping of the enriched canonical and Pan-Immune pathways overlapping between C1/C2 vs. C3-enriched and *KRAS-TP53* co-altered vs. *KRAS*-altered PDAC transcriptomes revealed significant conflation of biologic processes related to cellular heterogeneity and progenitor-like cancer stemness properties (Clusters A and F, respectively), as well as interconnectedness of pathways related to innate immune response (cluster G), epigenetic regulation and chromosomal maintenance (cluster B), and chromatin assembly (cluster D; **Fig 5B**).

**Figure 5:**
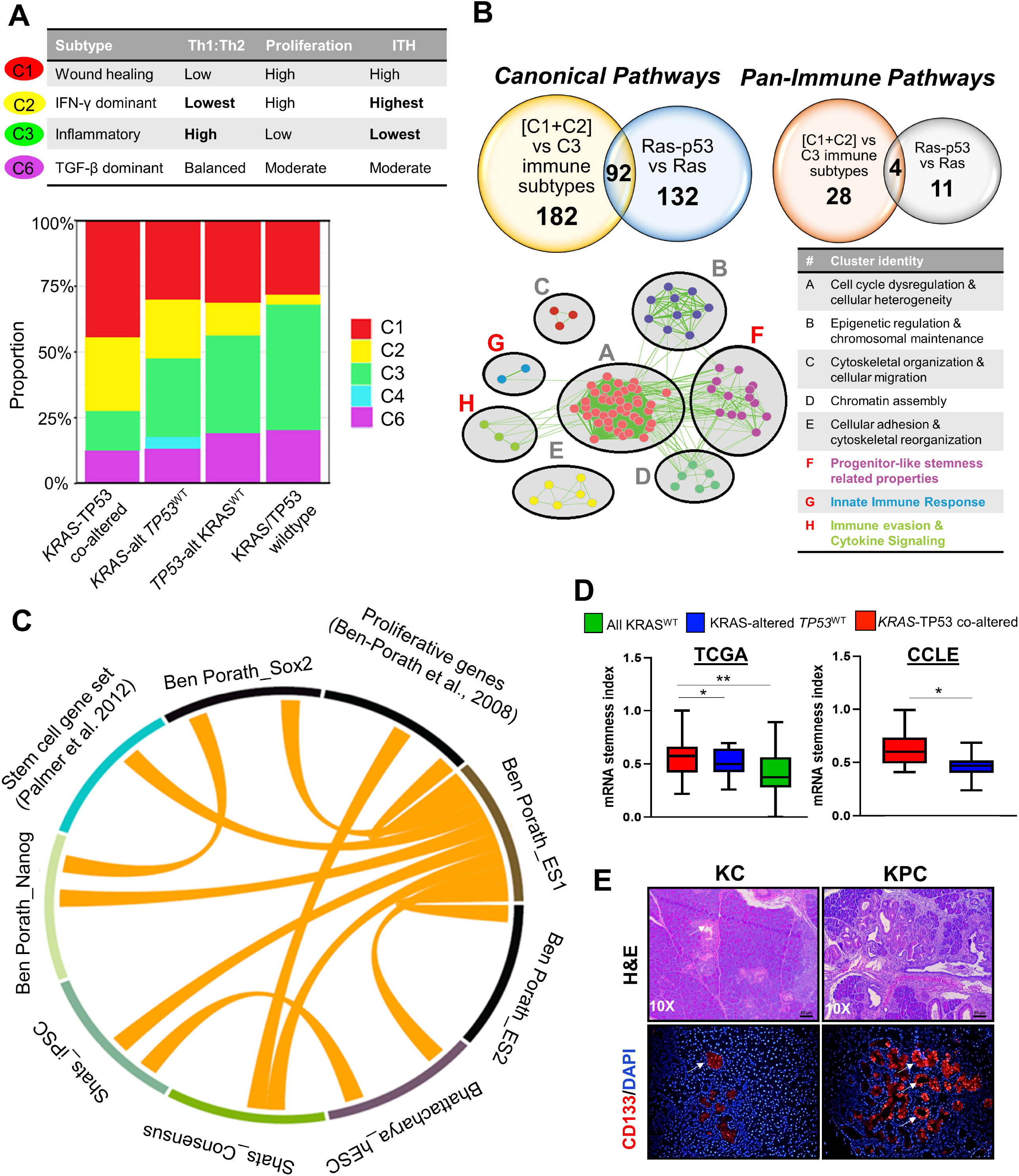
Consensus immune subtyping reveals conflation of intratumor heterogeneity with augmented stemness properties in *KRAS-TP53* co-altered PDAC. **(A)** Consensus immune subtype classification for C1, C2, C3, and C6 are shown in the table, denoting characteristics related to Th1:Th2 ratio, proliferation, and intra-tumor heterogeneity (ITH). Stacked histogram depicts relative frequency of these immune subtypes in TCGA-PAAD samples, stratified by *KRAS-TP53* co-altered, *KRAS*-altered *TP53*^WT^, *TP53*-altered *KRAS*^WT^, and *KRAS*-*TP53*-wildtype genomic status; **(B)** Venn diagrams showing overlap of pathways differentially expressed between C1+C2vs.C3 and *KRAS-TP53* vs. *KRAS*-altered/*TP53*^WT^ transcriptomes (FDR-adj P<0.1), for both canonical and Pan-Immune pathways. These 96 pathways were then mapped to an enrichment map showing interconnectedness of clusters A-H with identities tabulated in the adjacent table; **(C)** The differentially expressed *KRAS-TP53* co-altered TCGA-PAAD transcriptome overlaps with 9 gene sets representing stemness pathways enriched in cancer and embryonic cells, depicted with a Circos plot where links between two pathways suggest significant overlap (Jaccard index>0.05) between the differentially expressed genes (with FDR-adj P<0.2) involved in the respective pathways; **(D)** mRNA stemness index was calculated using TCGAanalyze_Stemness from the TCGAbiolinks package and compared between TCGA-PAAD *KRAS-TP53* co-altered, *KRAS* -altered *TP53*^WT^, and *KRAS*^WT^ transcriptomes (*left*) and *KRAS-TP53* co-altered vs. *KRAS*-altered *TP53*^WT^ human PDAC cells line transcriptomes from the Cancer Cell Line Encyclopedia/Depmap portal; **(E)** Representative sections showing H&E staining (*top*) and immunofluorescence staining for CD133/Prom1 (*bottom*) in tissue sections derived from KPC (age 5-6 months) and KC (age 12-14 months) genetic mouse models. \**p*<0.05; \*\**p*<0.01

Since tumor stemness characteristics have recently been linked to immune exclusion [26], we further dissected this association between *KRAS-TP53* co-alteration and stemness. We observed substantial overlap between the differentially expressed *KRAS-TP53* co-altered TCGA-PAAD transcriptome and 9 gene signatures associated with stemness in cancer and embryonic cells (**Fig 5C**); these sets spanned 2,613 unique genes (**Table S3**). Moreover, while several stemness-associated genes derived from the *ImSig* proliferation dataset were significantly overexpressed in TCGA *KRAS-TP53* co-altered vs. *KRAS*-altered/*TP53*^WT^ PDAC— *SHCBP1, OIP5, MAD2L1, STMN1, DLGAP5*, and *HMGB3* (**Fig S7A**)*—*pathway-level enrichment of proliferation/stemness-related *ImSig* genes was more strongly correlated with squamous vs. non-squamous, rather than *KRAS-TP53* genomic, stratification (**Fig S7B**).

The acquisition of a de-differentiated oncogenic phenotype and of progenitor, stem cell-like features—measured by a metric recently described as the cancer stemness index (SI)—has been linked to ITH and immune exclusion in a pan-cancer TCGA analysis [26]. We observed significantly higher mRNA-based SI (mRNAsi) in *KRAS-TP53* co-altered PDAC compared to *KRAS*-altered/*TP53*^WT^ and *KRAS*^WT^ tumors (mean 0.56±0.17 vs. 0.51±0.13 vs. 0.43±0.23, P=0.006) in the TCGA dataset, as well in *KRAS-TP53* co-altered vs. *KRAS*-altered/*TP53*^WT^ CCLE cell lines (mean 0.64±0.16 vs. 0.46±0.14, P=0.01) (**Fig 5D**). The latter observation was confirmed by increased expression of stemness markers CD133, CXCR4, ALDH1, and EpCAM in five *KRAS-TP53* co-altered (i.e., ASPC1, BxPC3, MiaPaCa-2, SW1990, and Panc1) vs.

*KRAS*-altered/*TP53*^WT^ (Hs766T) human PDAC cell lines via flow cytometric analysis (**Fig S7C**). To determine if the disproportionate stemness features in *KRAS-TP53* co-altered PDAC were driven by tumor-cells or other constituents of the PDAC TME, we observed significant overexpression of several stemness-associated genes (i.e., *Prom1, Stmn1, Muc1, Itga6, Itgb1, Alcam, Cd24a*, and *Hmgb3*) in *Kras-Trp53* co-altered vs. *Kras*-altered/*Trp53*^WT^ tumor-cell clusters via scRNAseq (**Fig S7D**). These transcriptomic differences were then confirmed by significantly increased immunoreactivity of CD133/Prom1 staining in the epithelial compartment of tumor tissue derived from KPC compared with KC PDAC GEMMs (**Fig 5E**). These data suggest that the disproportionate enrichment of immune subtypes associated with ITH and progenitor-like stemness properties in *KRAS-TP53* co-altered PDAC may underlie immune exclusion and worse oncologic outcomes observed in these patients.

### Development of an immunoregulatory transcriptional program associated with KRAS-TP53 co-alteration in PDAC

We combined 20 genes differentially overexpressed in *KRAS-TP53* co-altered PDAC— associated with squamous transdifferentiation, innate immunoregulation, adaptive immune evasion, inflammasome machinery, and stemness features (**Table S6**)—into a gene signature to explore its clinical relevance with respect to immune exclusion, chemoresistance, and survival in PDAC patients. By applying immune deconvolution analysis in the TCGA dataset, overexpression of this 20-gene signature was associated with significant enrichment of macrophage and neutrophil populations, checkpoint molecules, and inflammation-promoting pathways, as well as downregulation of helper/effector T-cell populations (**Fig 6A**). Given these findings, we termed this 20-gene signature the “*KRAS-TP53* co-altered immunoregulatory program” or *KRAS-TP53* IRP.

**Figure 6:**
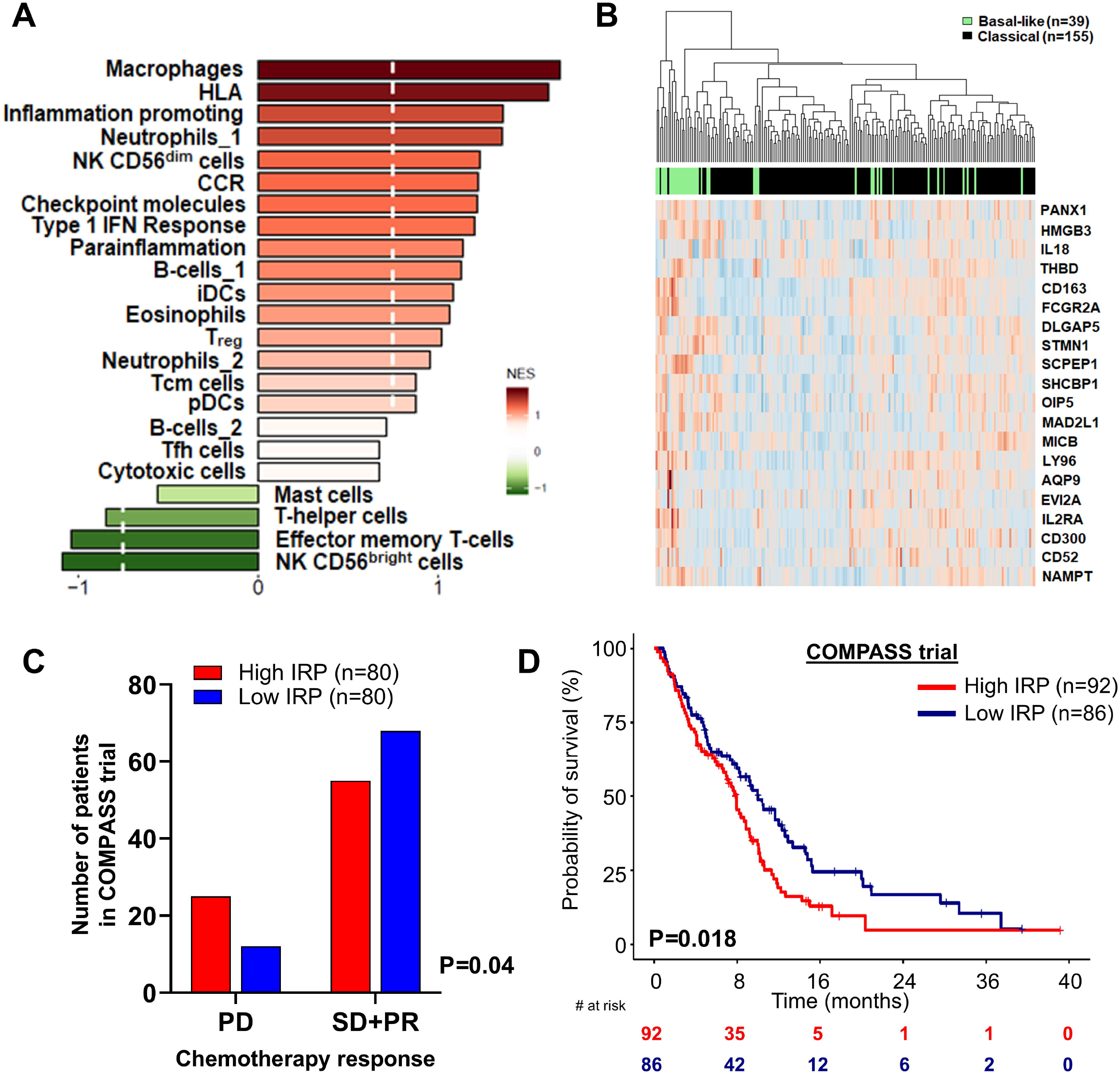
Development of an immunoregulatory transcriptional program associated with *KRAS-TP53* co-alteration in PDAC. **(A)** Curated immune and inflammation-focused gene set enrichment analysis showing immune populations and pathways differentially expressed in *KRAS-TP53* immunoregulatory program (IRP) “high” versus “low” samples in TCGA-PAAD dataset. Broken white line indicates FDR-adj P<0.05; **(B)** Heatmap depicting that expression of the 20 genes comprising the *KRAS-TP53* IRP gene signature distinguish samples classified as basal-like (green) and classical (black) molecular subtypes in the COMPASS trial dataset (n=195); **(C)** Within the subset of patients in the COMPASS trial dataset with annotated chemotherapy response information (n=160), the histogram represents the number of patients demonstrating either progressive disease (PD) or stable disease (SD)/partial response (PR), stratified by *KRAS-TP53* IRP high (*red*) or low (*blue*) gene signatures; **(D)** Overall survival curve comparing Kaplan-Meier estimates between patients with high and low *KRAS-TP53* IRP gene signatures from the COMPASS trial dataset. The IRP score corresponds to the number of genes (of 20) per sample whose expression is >median expression of the individual gene in the COMPASS dataset. Subjects are dichotomized into a “high” (score>10) and “low” IRP (score<10) with samples with a score=10 excluded. The adjoining table details number of patients at risk at designated time points.

Next, utilizing the COMPASS trial dataset which included 195 advanced PDAC patients undergoing RNA sequencing—of which 160 patients had annotated RECIST response rate data following multi-agent chemotherapy [34]—we observed that the *KRAS-TP53* IRP was strongly correlated with basal-like molecular subtype (**Fig 6B, Fig S8A**), previously associated with chemoresistance and worse survival [34]. As such, in the COMPASS dataset, patients harboring high *KRAS-TP53* IRP-expressing tumors (n=80) were significantly more likely to demonstrate progressive disease, whereas low *KRAS-TP53* IRP-expressing tumors (n=80) were more likely to demonstrate stable disease or partial response to chemotherapy (P=0.04). Notwithstanding the lack of additional clinically annotated PDAC datasets for validation, we observed disproportionate enrichment of high *KRAS-TP53* IRP expression in five independent gene sets associated with chemoresistance in ovarian [35], genitourinary [36], lung [37], and breast [38] malignancies (**Fig S8B**).

Finally, high *KRAS-TP53* IRP-expressing PDAC was associated with significantly worse overall survival compared with low *KRAS-TP53* IRP-expressing PDAC in the COMPASS dataset (median 7.9 vs. 10.0 months, P=0.018) (**Fig 6D**). The prognostic disadvantage of high *KRAS-TP53* IRP expression in PDAC was validated in both the ICGC (median 14.6 vs. 22.5 months, P=0.048) and TCGA (median 19.6 vs. 23.1 months, P=0.067) datasets (**Fig S8C**). Collectively, these data highlight the clinical relevance of an immunoregulatory gene signature which incorporates myriad elements of molecular determinants distinctive to the *KRAS-TP53* co-altered PDAC transcriptome.

## DISCUSSION

Given that co-occurrent *KRAS-TP53* alterations are observed in nearly two-thirds of all PDAC patients and drives aggressive pro-metastatic phenotypes [10], the present study is the first to demonstrate that *KRAS-TP53* co-alteration is associated with worse survival compared to mutually exclusive *KRAS* or *TP53* alterations in heterogeneous populations of PDAC patients. We leverage this novel observation to decipher its unique molecular determinants and their association with immune regulation using an integrative data-driven approach with *in vitro* and *in vivo* validation in preclinical models. These molecular determinants are coalesced into a *KRAS-TP53* “immunoregulatory program” (IRP)—encompassing both cancer cell-autonomous and non-autonomous factors—which is associated with chemoresistance and worse survival in PDAC patients. Individually, elements of this IRP provide a biologic map of the underexplored genotype-immunophenotype chasm in PDAC and uncover novel targets for immunomodulatory therapies that could overcome therapeutic resistance in this disease.

While the independent roles of both oncogenic *KRAS* activation and *TP53* mutations in establishing pro-inflammatory signaling and activation of immunosuppressive cells are well established [53, 54], how *KRAS-TP53 co-alteration* encode unique transcriptional programs to govern immunologic remodeling of the PDAC TME is incompletely understood. As such, our data expand on recent insights suggesting that *KRAS-TP53* co-alteration disproportionately controls the trafficking of innate immune populations into established tumors [55] and that *TP53* missense mutations in PDAC orchestrate T-cell exclusion [11], suggesting that innate immunoregulatory ecosystems in PDAC may be dependent on its genomic context. Intriguingly, our data not only uncover potential cancer cell-autonomous transcriptional programs encoded by *KRAS-TP53* co-alteration that orchestrate innate immune cell trafficking, but also implicate granulocyte-derived inflammasome and *TNF* signaling as novel *functional* regulators of the dynamic immune-tumor-stromal crosstalk in the TME that sustain innate immunoregulatory signaling in *KRAS-TP53* co-altered tumor cells. Indeed, emerging evidence highlights the paradoxical pro-tumorigenic roles of both inflammasome and *TNF* signaling in cancer initiation and progression [56, 57]. Furthermore, our lineage trajectory reconstruction in single-cell intratumoral gMDSC transcriptomes identify distinct granulocytic developmental fates associated with *Tnf* signaling, and provides a molecular roadmap for interception strategies aimed at disrupting gMDSC compartment-specific TNF signaling in PDAC. Taken together, these data suggest that cellular lineage-restricted and genomic context-dependent disruption of *TNF* signaling and inflammasome activation may emerge as attractive strategies to reprogram the immune-excluded TME and overcome immunotherapeutic resistance in PDAC.

Although recent evidence has implicated squamous trans-differentiation as a master regulator of stromal inflammation and innate immune infiltration in PDAC, the upstream mechanisms that induce ΔNp63/TP63-driven squamous transdifferentiation have remained elusive [45]. Our data suggest that *KRAS-TP53* co-alteration may be one of the incipient genomic events that induces such squamous trans-differentiation and its deleterious downstream repercussions. More provocatively, since consensus immune subtyping [18] reveals a novel association between *KRAS-TP53* co-alteration and increased intra-tumor heterogeneity, it is plausible that the spatial and temporal evolution of treatment-resistant squamous subclones [58] during therapeutic trajectories in *KRAS-TP53* co-altered PDAC might sculpt progressively tolerogenic immune ecosystems and dictate its chemoresistant microenvironment. It is possible, therefore, that therapeutic strategies targeting squamous transdifferentiation-defining signaling pathways (e.g., IL-1 signaling) in PDAC may overcome these elements of its aggressive biology. As such, data from the currently accruing Precision Promise™ adaptive design phase 2/3 trial (NCT04229004) investigating inhibitors of IL-1β (canakinumab, Novartis) and IL-1RAP (CAN04, Cantargia) in metastatic PDAC patients are eagerly awaited.

Given the link between mRNA stemness index and immune exclusion, reduced PD-L1 expression, and pro-metastatic proclivity in pan-cancer analyses [26], the association between *KRAS-TP53* co-alteration and elevated stemness features in the present study provide additional insight into its therapeutically resistant and immunologically “cold” biology. More importantly, the discovery of novel stemness-related transcripts that may impart progenitor-like properties in *KRAS-TP53* co-altered PDAC—e.g., HMGB3, targeting which overcomes platinum resistance in ovarian cancer via an ATR/CHK1-mediated mechanism [59]—could be leveraged into novel therapies aimed at overcoming immune exclusion and therapeutic resistance in PDAC.

This IRP, curating diverse elements of tumor-permissive molecular crosstalk engineered by *KRAS-TP53* co-alteration, adds to an expanding compendium of clinically relevant molecular taxonomy that highlights the clinical and phenotypic heterogeneity observed in PDAC patients. These data also attempt to address the critically underexplored genotype-phenotype chasm in PDAC, suggesting that high-risk *KRAS-TP53* co-altered PDAC genotypes elicit distinct and non-redundant transcriptional programs to amplify innate immunoregulatory signaling and thwart effector immune responses. Our data uncover novel molecular susceptibilities that underpin these transcriptional networks which could be exploited to not only develop biomarkers predicting therapeutic response and resistance, but also treatment strategies to overcome immune exclusion and sensitize PDAC to chemo- and/or immunotherapy. Ultimately, these data emphasize that a multi-pronged approach will be needed to effectively subvert immune tolerance, unleash anti-tumor immunity, and revolutionize treatment for patients with PDAC.

## ACKNOWLEDGEMENTS

This work was supported by KL2 career development grant from Miami CTSI under NIH Award UL1TR002736, Stanley Glaser Foundation, American College of Surgeons Franklin Martin Career Development Award, and Association for Academic Surgery Joel J. Roslyn Faculty Award (to J. Datta); NIH R01 CA161976 (to N.B. Merchant); and NCI/NIH Award P30CA240139 (to J. Datta and N.B. Merchant).

## AUTHOR CONTRIBUTIONS

JD designed the study; JD, NBM provided funding; JD, YB, AB, IDCS, NUD, LLC, SM, SS, CR, XS, XD, AC, PS, ARD performed the experiments; JD, AP, PJH, NSN, JMW, JJK, XC, KVK, NBM provided access to clinical/murine samples; JD, YB, AB, CR wrote the manuscript; all authors reviewed/edited the manuscript.

## DATA AVAILABILITY

The authors declare that the data supporting the findings of this study, and information for retrieval of all external sources of data, are available within the manuscript. The detailed mutational information of tumor samples in the UMiami cohort (n=245) was collated from commercial vendors in the UMiami Patient Atlas, and putative oncogenic mutations are provided in **Table S1**.

