## Supplementary Materials (Method, Figures, Tables S6-S7) for "Distinct Mechanisms of Innate and Adaptive Immune Regulation Underlie Poor Oncologic Outcomes Associated with *KRAS-TP53* Co-Alteration in Pancreatic Cancer"

A. Supplementary Methods……………………………………………1-5

B. Supplementary References………………………………………...6-7

C. Supplementary Figures with Legends.....…………………………..8-16

D. Supplementary Table S1-S5 Legends……………...………………..17

F. Supplementary Table S6………………………………………………18

G. Supplementary Table S7…………………..…...………………………19

### A. Supplementary Methods

*University of Miami Clinical Cohort*

After obtaining Institutional Review Board approval for the study, patients with metastatic PDAC who underwent treatment at the University of Miami/Sylvester Comprehensive Cancer Center between 7/2015 and 12/2019 and had available NGS testing using either of two commercially available platforms (Caris™ and FoundationOne™) were identified from the UMiami Patient Atlas™, a molecular repository harmonizing health system-wide NGS testing. Patients with microsatellite-stable (MSS) tumors and alterations in *KRAS*, *TP53*, or both (n=245) were retrieved and annotated by the OncoKB knowledgebase [1] to designate putative driver or structural alterations. Clinical annotation of these NGS samples were performed by retrospective review of the electronic medical records to collect information on patient age, gender, date of birth, date of diagnosis, stage at presentation, primary tumor location in pancreas, whether primary tumor had been previously resected or if patient presented with synchronous metastatic disease, sites of metastasis, chemotherapy exposure prior to submission for NGS, and chemotherapy regimen (**Table S1**). The choice of chemotherapy regimen and dosage was selected at the discretion of the medical oncologist. We confirmed that NGS testing for all included patients was performed prior to any patient's enrollment in a clinical trial and receipt of investigational therapy.

The date of last follow-up and/or date of death as well as vital status were accrued. The primary endpoint was overall survival (OS), defined as duration between date of diagnosis and date of death or last follow-up. We were not able to accurately capture recurrence/progression information on a large number of patients due to interval imaging and treatments at sites outside of our tertiary referral cancer center.

*Statistical Analysis of Clinical Data*

Descriptive statistics were calculated for patients' characteristics using median, interquartile range (IQR), frequencies, and 95% confidence intervals (95% CIs). All tests were two-sided and statistical significance was considered when $p \leq 0.05$. Given the absence of distinguishing demographic and clinicopathologic characteristics between patients in the *KRAS-TP53* co-altered, *KRAS*-altered $TP53^{WT}$, and *TP53*-altered $KRAS^{WT}$ cohorts (**Table S1**), we compared OS using log-rank tests with Kaplan-Meier estimates. Survival analysis was performed using R package "survival" (https://cran.r-project.org/web/packages/survival) and visualized using "survminer" (https://cran.r-project.org/web/packages/survminer/).

*Public Data Retrieval*

TCGA Pan-Cancer gene-level non-silent somatic mutation (v0.2.8) data and TCGA PAAD gene expression log2-transformed RSEM-normalized data matrices were retrieved from Xena [2]. Previously identified PAAD molecular subtype [3] information was extracted using TCGAbiolinks [4]. TCGA-PAAD immune subtyping was performed by stratifying into consensus molecular subtypes C1-C6, as described previously [5], using the TCGAbiolinks tool [4]. RNA sequencing data (bam files) and survival information from the COMPASS trial dataset were obtained from EGA data set EGAD00001006081 under a previous study [6]. Gene expression count matrix was generated using featureCounts [7] on the retrieved bam files for COMPASS data. ICGC-PACA-CA data set including RNAseq (FPKM-normalized) and survival information was downloaded from ICGC Data Portal (https://dcc.icgc.org/). Single-cell RNA sequencing datasets "Late KIC" (GEO #GSM3577884) and "Late KPfC" (GEO #GSM3577885) were downloaded from the Gene Expression Omnibus (GEO) database [8]. Panc02 single-cell RNA sequencing dataset was retrieved from the Sequence Read Archive (SRA) NCBI repository (#SRX7873760) [9].

*mRNA stemness index assignment*

Stemness scores were calculated as previously described [10]. Briefly, to calculate the stemness scores based on mRNA expression, we built a predictive model using one-class logistic regression (OCLR) [11] on the pluripotent stem cell samples (ESC and iPSC) from the Progenitor Cell Biology Consortium (PCBC) dataset [12, 13]. For mRNA expression-based signatures, to ensure compatibility with the TCGA PAAD cohort, we first mapped the gene names from Ensembl IDs to Human Genome Organization (HUGO), dropping any genes that had no such mapping. The resulting training matrix contained 12309 mRNA expression values measured across all available PCBC samples. RNA-seq raw counts of TCGA-PAAD and CCLE human PDAC cell line samples aligned to the hg19 reference genome were downloaded from their respective archives, normalized, and filtered using the R/Bioconductor package TCGAbiolinks14 version 2.9.5 using GDCquery(), GDCdownload(), and GDCprepare() functions for tumor types (level 3, and platform "IlluminaHiSeq_RNASeqV2"), as well as using data.type as "Gene expression quantification" and file.type as "results". Then tumor matrix was then normalized using within-lane normalization to adjust for GC-content effect on read counts and upper-quartile between-lane normalization for distributional differences between lanes by applying the TCGAanalyze_Normalization() function adopting the EDASeq protocol. To calculate mRNA based stemness index (mRNASi) we used the function TCGAanalyze_Stemness from the package TCGAbiolinks and following our previously-described workflow, with "stemSig" argument set to HipSci_532_iPs-stemsig.tsv.[4]

*Single-cell RNA sequencing cluster identification using Seurat*

The Seurat pipeline (version 3.2) was used for cluster identification in Panc02, "late KIC" and "late KPfC" scRNAseq datasets, retrieved as described previously [14]. Data were read into R Studio (version 4.0.2) as a count matrix, scaled by a size factor of 10,000, and log transformed. Gene expression cutoffs were set at a minimum of 200 and a maximum of 7500 genes in more than three cells for each dataset. In addition, cells with a percentage of mitochondrial genome (referred to as percent.mt) greater than 8% were removed. KIC and KPfC datasets were integrated into a single-combined analysis. For tSNE/UMAP projections and clustering analysis, we computed 30 principal components. Markers for each cluster identified by Seurat were determined using the "FindAllMarkers" function. Clustering via the "FindClusters" function was generated, and subsequent cluster labeling was performed based on the top 50 genes differentially upregulated per cluster. DE genes (FDR-adjusted P<0.05) in the tumor-cell cluster of KPfC (*Kras-Tp53* co-altered) vs. KIC (*Kras*-altered/*Tp53*$^{WT}$) datasets were identified using the "limma" package, and visualized using volcano plots and Cytoscape network analysis.

*Single-cell lineage trajectory reconstruction*

Using the R package Monocle (version 2.8.0) [15], a differentiation hierarchy within the Panc02 granulocytic myeloid compartment was reconstructed. After removal of contaminating cell types, the granulocyte cluster was selected for subsequent analysis and re-clustered to explore additional heterogeneity within this cellular compartment. Using unbiased clustering, the top 20 unique genes per cluster were used to order cells along a pseudo-temporal trajectory.

The cell clusters obtained in Monocle were exported and utilized as input to analyze single cell lineage using the RNA velocity tool velocyto [16] with default parameters. Normalization and clustering were performed using PAGODA2 [17].

*Orthotopic murine PDAC model generation and in vivo studies*

All animal experiments were performed in accordance with the NIH animal use guideline and protocol approved by the Institutional Animal Care and Use Committee (IACUC) at the University of Miami. C57BL/6 female mice (6 weeks age) were purchased from Jackson Laboratory (Bar Harbor, ME) and housed at the institutional animal facility under constant temperature and humidity, on a 12-hour light and dark cycle, with available standard food and filtered water. For orthotopic model generation, C57BL/6 mice were anesthetized with Ketamine:Xylazine (10:1) in sterile 0.9% NaCl saline. The abdomen was then wiped with iodine-based solution followed by alcohol wipes. The mice were placed on a clean sterile pad onto a warming pad. A paramedian incision was made in the abdomen using scissors and the pancreas and spleen were externalized with assistance of curved forceps. Then, 10 µL of cell suspension containing 200,000 *LSL-KrasG12D*$^{/+}$*; LSL-Trp53R172H*$^{/+}$*; Pdx1*$^{Cre}$ (KPC-6694c2) or *LSL-KrasG12D*$^{/+}$*; Pdx1*$^{Cre}$ (KC-PDA4313) tumor cells in Matrigel® were injected directly into the pancreas using a Hamilton™ microliter syringe. The pancreas and spleen were then returned to the intra-abdominal cavity in the anatomical position and the peritoneum sutured with Vicryl 5-0 suture and the skin closed with skin staples. Immediately after surgery, 300µL of sterile saline was injected subcutaneous for fluid maintenance and buprenorphine 0.05-0.1mg/kg was given subcutaneously for pain management. Mice were continuously monitored post-operatively and staples were removed two weeks post-surgery. Mice were sacrificed when the tumor volume for mice in each group (i.e., KPC, KC) achieved 1 $cm^3$ (albeit at different time points), and tumor and spleen samples were collected for subsequent analysis.

*Tissue collection and cell isolation*

**Bone marrow**. Tumor-naïve mice bone marrow (BM) cells were flushed from mouse tibia and femurs using a 28G needle and plastic syringe and then kept in RPMI 1640 (Corning, 10-041-CV3J), 10% FBS (VWR). BM cells were centrifuged at 350g, 4°C for 5 min. Cells were incubated for 3-4 min at room temperature in 2 mL red blood cell (RBC) lysis buffer. Cells were quenched with 20 ml complete medium, and centrifuged at 350g, 4°C for 5 min.

**Spleen**. Spleens from KC and KPC tumor-bearing mice were minced using a syringe filter, and passed through 100 µm mesh filters to obtain spleen single-cell suspensions. These suspensions were then processed using RBC lysis buffer. Ly6G$^+$ cells were isolated from fresh suspensions using the Myeloid-Derived Suppressor Cell Isolation Kit, LS Columns, and QuadroMACS™ Separator (Miltenyi Biotech).

**Pancreas/tumor**. Whole pancreata harvested from C57BL/6 mice were digested enzymatically utilizing a solution containing 0.6 mg/ml of collagenase P (Roche), 0.8 mg/ml Collagenase V (Sigma Aldrich), 0.6 mg/ml soybean trypsin inhibitor (Sigma Aldrich), and 1800 U/ml DNase I (Thermo Scientific) in RPMI medium for 20-30 minutes at 37°C. Samples were then washed and subjected to single cell dissociation using 40 µm smash strainers. For isolation of individual cell populations for functional inflammasome assay (see below), whole tumor single-cell suspensions were stained with EpCAM, PDPN, Ly6G, and Ly6C markers (**Table S7**) and individual cell populations sorted using CytoFLEX S FACS sorter (Beckman Coulter).

*In vitro generation of gMDSCs and migration assay*

BM cells were collected as described above, and cells were cultured with RPMI and 10% FBS and treated with 20ng/ml recombinant murine GM-CSF and IL6 (R&D systems) on day 1 and on day 3. We then confirmed that the majority of cells obtained through this conditioning were $CD11b^{+}F4/80^{-}Ly6G^{+}Ly6C^{dim}$ (i.e., gMDSCs). gMDSCs were then placed on top of the filter membrane in a transwell insert (8μm pore size), and 48h-serum free-conditioned media from KC, KPC, and KPC-*Cxc1*$^{KO}$ [18] tumor cells was added at the bottom of the lower chamber, serving as chemo-attractant source. Six hours after, MDSCs that migrated to the lower chamber were detected with LUNA™ Automated cell counter (Logos Biosystems).

*Caspase-1 luminescence assay for functional inflammasome activity*

Caspase-1 activity was measured using the Caspase-Glo® 1 Inflammasome Assay (Promega, #G9951) as per the manufacturer's protocol. Freshly isolated $EpCAM^{+}$, $PDPN^{+}$, $F4/80^{-}Ly6G^{+}$ and $F4/80^{+}$ cells from whole tumor suspensions from KC and KPC orthotopic tumors were seeded in a 96-well plate and treated with caspase-1 substrate (Z-WEHD-aminoluciferin) conjugated to a fluorophore, followed by incubation at 37°C for two hours. The luminescence was measured using SpectraMax® iD5 (Molecular Devices). All positive and negative controls were included as per the manufacturer's protocol. All measurements were carried out in biological triplicates.

*Multiplex cytokine array*

Cell lysates from human PDAC Ras-alone (Hs766t, KC-PDA4313), and human PDAC Ras-p53 co-altered cell lines (MiaPaca2, Capan1, Panc02.03, and KPC-6694c2 and/or K8484) were analyzed using the Proteome Profiler Human Cytokine Array Kit (#ARY005B, R&D Systems, Minneapolis, MN, USA) according to the manufacturer's instructions.

*Transfection of TP53 constructs in HPNE-KRAS$^{G12D}$ cells and Western blotting*

KC and KPC tumors, as well as HPNE-*KRAS*$^{G12D}$ cells transfected with either mutant *TP53*$^{R175H}$ or *TP53*$^{WT}$ cDNA constructs, were homogenized (Omni™ Tissue Homogenizer) in 0.5 ml of RIPA lysis buffer (20-188, Millipore®) and sonicated for 2 minutes on ice. Tissue lysates were centrifuged for 10 min at 15000 rpm at 4°C. Supernatants were then collected, quantified with BCA assay (23227; ThermoFisher Scientific). Standard procedures were used for western blotting, as extensively described previously [19-21]. Primary antibody used were Vinculin (13901; Cell Signaling), and p40DeltaNp63 (ab203926; abcam). Proteins were detected using HRP-conjugated secondary antibodies (Jackson ImmunoResearch Laboratories).

*Flow Cytometry*

Fresh tumor and spleen single-cell suspensions were thawed, washed, and incubated with FcR blocking reagent (Miltenyi Biotec), and subsequently stained with fluorescently conjugated antibodies (**Table S7**). Ghost Red Dye 780 (TONBO biosciences) live/dead cell discrimination was performed as per manufacturer's protocol and cells were fixed with 1% formaldehyde solution (Thermo Fisher). Flow cytometry data acquisition was performed on CytoFLEX S (Beckman Coulter) and analyzed using FlowJo v10 software (BD Life Sciences). For viSNE plot generation, flow cytometry data were analyzed using Cytobank software (Cytobank, Santa Clara, CA) with default parameters (iterations= 1000, perplexity= 30, θ= 0.5). All samples were derived from the same viSNE run by combining all individual flow cytometry standard files into a single flow cytometry standard file using the concatenation tool. viSNE heat maps show the fluorescent intensity of each marker for each event. Scales on the heat maps were individually generated for each surface marker from low to high expression.

*Histologic Analysis*

Pancreatic tissues were fixed in 10% neutral-buffered formalin followed by 70% ethanol after 24 hours and embedded in paraffin. For hematoxylin and eosin (H&E) staining, sections were mounted on glass slides and deparaffinized in xylene followed by rehydration using alcohol gradient. Thereafter, slides were stained with hematoxylin solution for 3 minutes and eosin solution for 1 minute and washed in between with running tap water. For immunohistochemistry (IHC) staining, sections were mounted on glass slides, deparaffinized and rehydrated followed by antigen retrieval which involved incubating samples in citrate buffer (0.01 M, pH 6:0) and heating. Sections were then blocked using BlockAid™ (Thermo Fischer) to preclude non-specific binding. For IHC experiments, endogenous peroxidase activity was quenched by incubating with 3% peroxide for ten minutes. Samples were then incubated with the primary antibodies (**Table S7**) in a humidified chamber at 4°C overnight. On the following day, slides were washed and developed using VECTASTAIN R Elite ABC HRP based kit (Vector) as per the manufacturer's protocol with diaminobenzidine (DAB) as the chromogen. Tissue sections were counterstained with Mayer's hematoxylin, mounted, and imaged.

For immunofluorescence staining (IF), after blocking, slides were incubated with primary antibody (CD133, ab19898, clone GR21491-1, abcam) polyclonal biotinylated secondary antibody and fluorophore-conjugated Streptavidin, followed by DAPI staining. Quantitative histological analysis was performed by sampling multiple random, non-overlapping 20x fields in each tissue section and quantified using ImageJ software, Fiji plugin (NIH) to measure the percent area positive stain per field (https://imagej.net/software/fiji/).

### C. Supplementary Figures

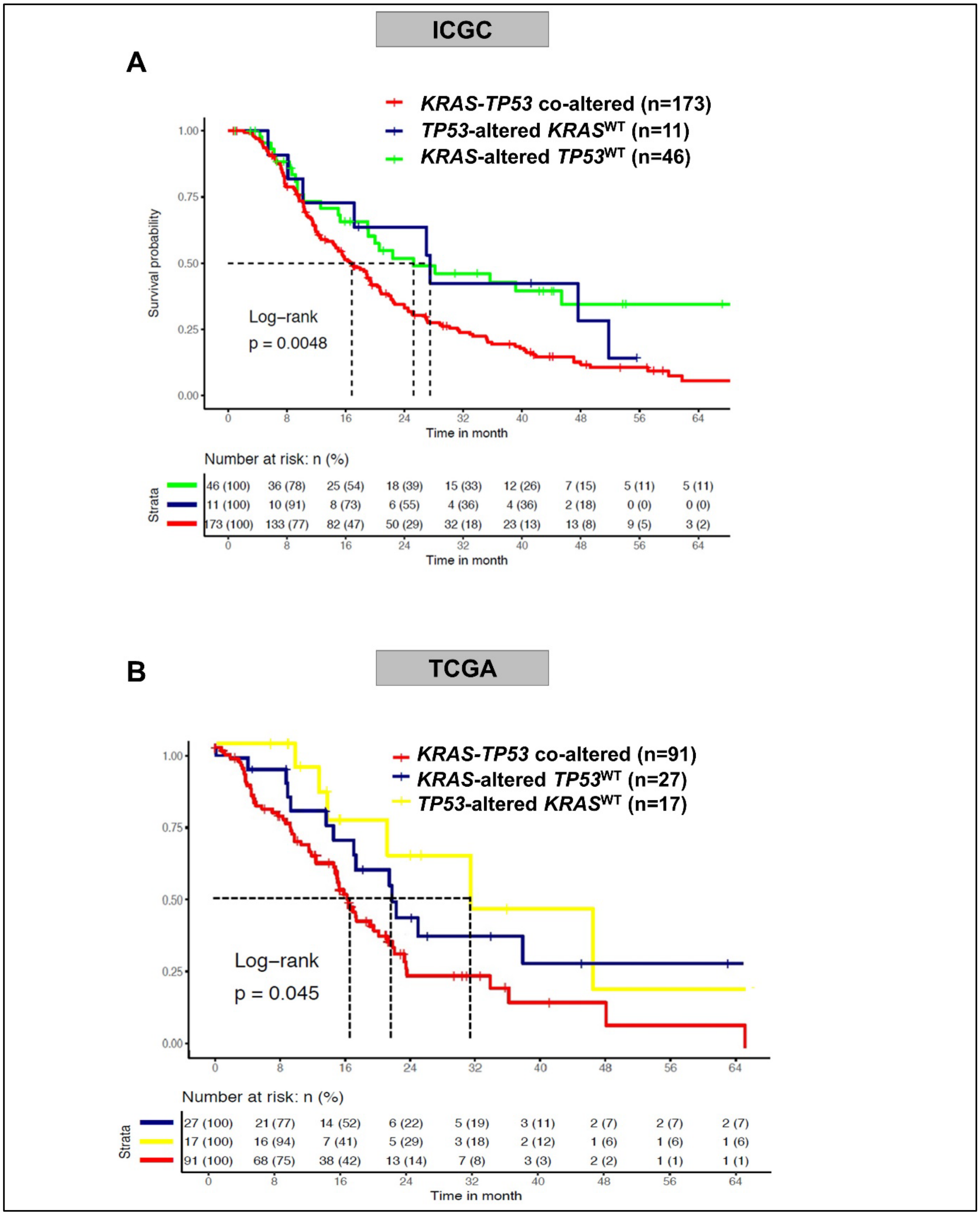


**Supplementary Figure S1**: Overall survival using Kaplan-Meier estimates in PDAC patients, stratified by *KRAS-TP53* co-altered, *KRAS*-altered *TP53*$^{WT}$ or *TP53*-altered *KRAS*$^{WT}$ mutational status in the: **(A)** ICGC, and **(B)** TCGA-PAAD datasets. Number at risk at each timepoint indicated in adjoining risk table in respective graphs.

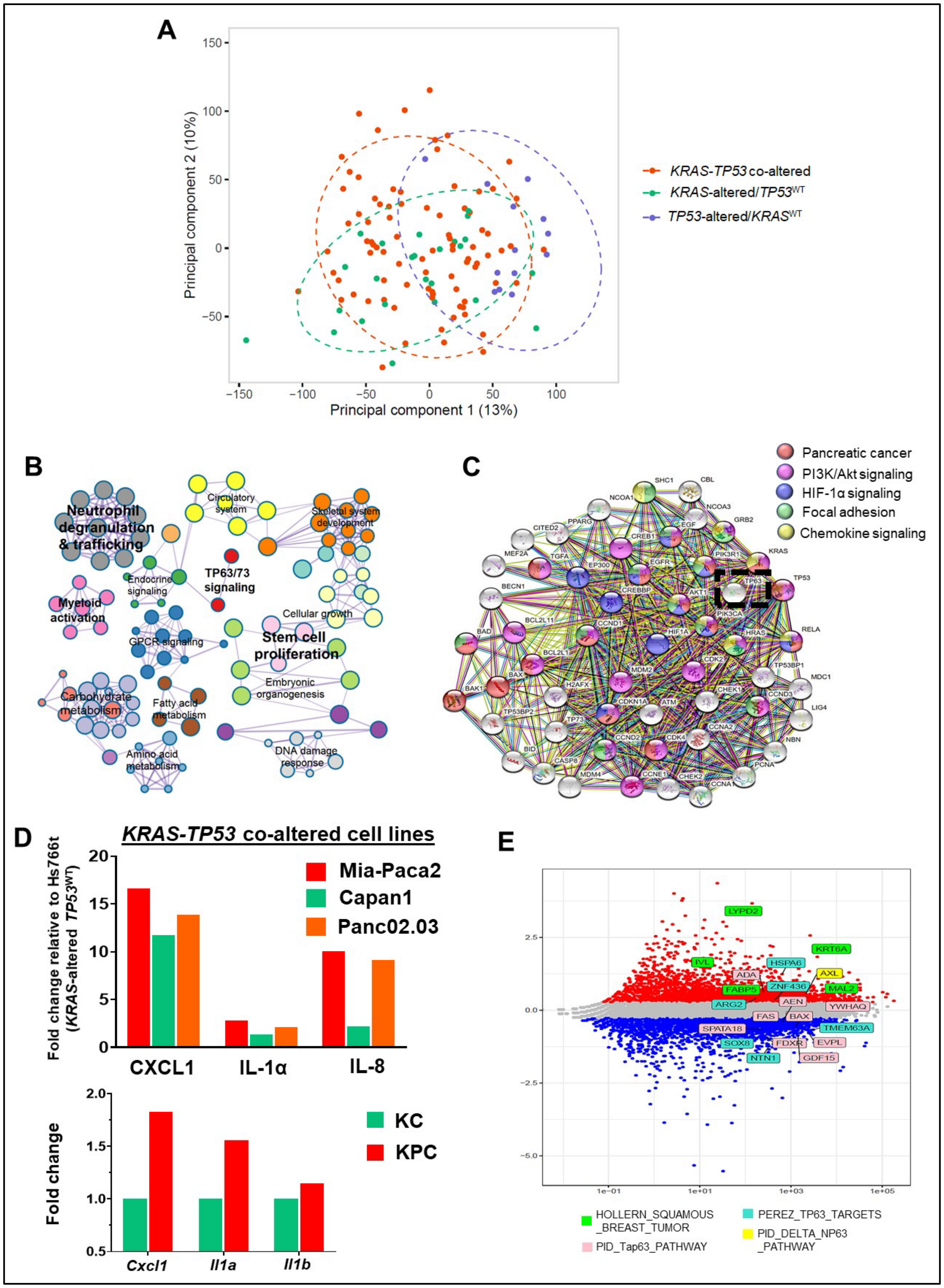


**Supplementary Figure S2: (A)** Principal component analysis of the transcriptional similarity of *KRAS-TP53* co-altered, *KRAS*-altered/*TP53*$^{WT}$ or *TP53*-altered/*KRAS*$^{WT}$ samples in the TCGA-PAAD dataset; **(B)** Cytoscape network analysis mapping the differentially expressed pathways in tumor transcriptomes of *KRAS-TP53* co-altered compared with *KRAS*-altered/*TP53*$^{WT}$ TCGA-PAAD samples; **(C)** protein-protein-interactome (PPI) of differentially expressed genes in *KRAS-TP53* co-altered vs. *KRAS*-altered/*TP53*$^{WT}$ PDAC (obtained from STRING portal) shows predicted biologic connectivity of TP63 (dashed box) to both KRAS and TP53. Other connected nodes in this PPI that are known to be involved in PDAC signaling, as well as in pathways (PI3K/Akt, HIF-1α, Focal Adhesion, Chemokine Signaling) upregulated in non-PDAC squamous tumors (anal, skin, etc.) are annotated in the adjoining legend; **(D)** Fold increase in secretion of cytokines implicated in defining squamous transdifferentiation in 3 *KRAS*-*TP53* co-altered human PDAC cell lines (Mia-Paca2, Capan1, Panc02.03) relative to *KRAS*-altered/*TP53*$^{WT}$ cell line Hs766t (top), and in KPC-6694c2 tumor cells compared with KC-PDA4313 cells (bottom), using multiplex cytokine arrays; **(E)** MA plot highlights differentially expressed genes in *KRAS*-*TP53* co-altered versus *KRAS*-altered/*TP53*$^{WT}$ TCGA-PAAD samples, previously implicated in pathways defining squamous lineage in non-PDAC solid tumors.

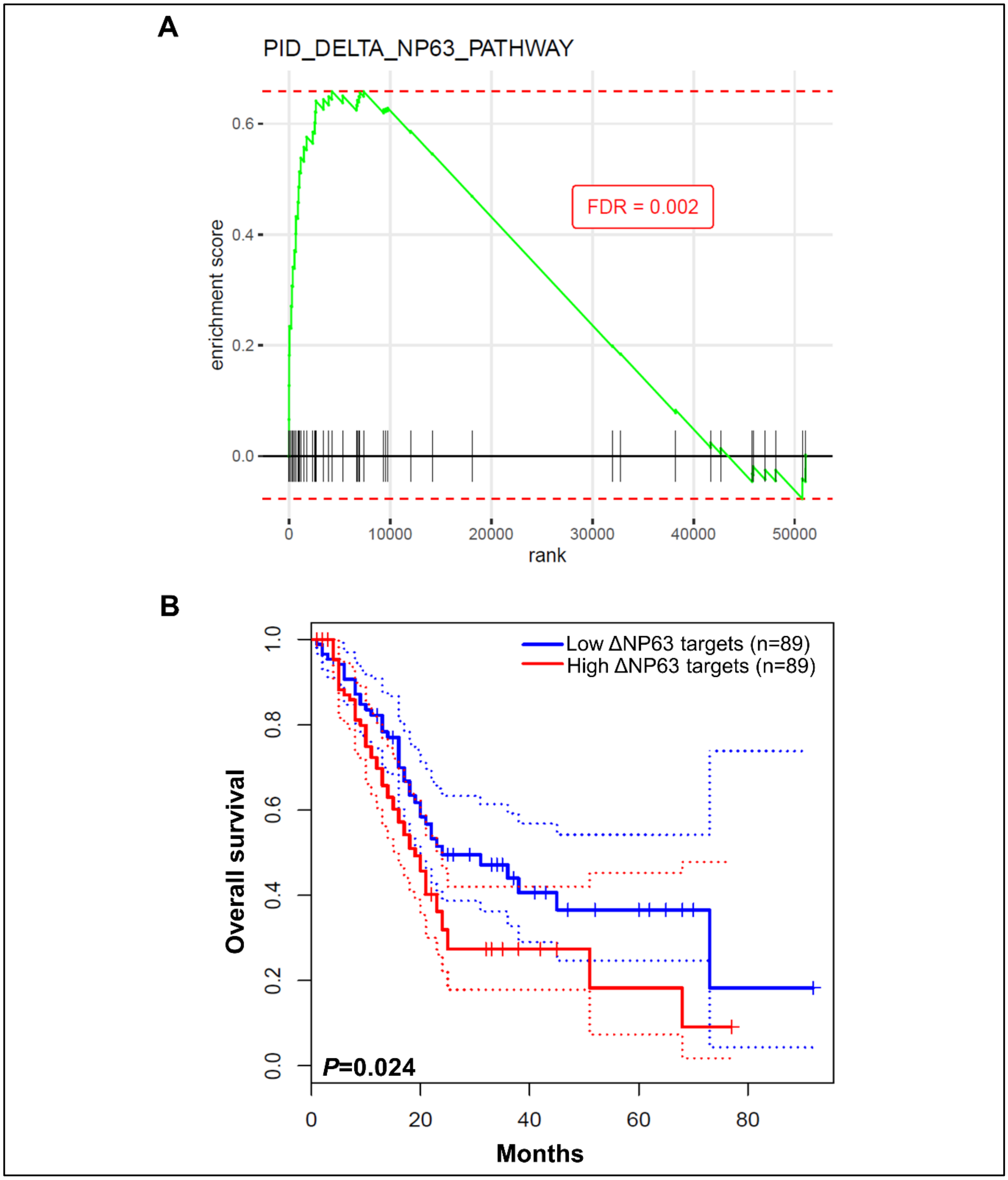


**Supplementary Figure S3**: **(A)** Enrichment plot depicting enrichment of transcripts in the PID_DELTA_NP63_PATHWAY in advanced PDAC patients in the COMPASS trial demonstrating progressive disease (chemoresistant) compared with stable disease or partial response (chemoresponsive; FDR: false discovery rate); **(B)** Kaplan-Meier overall survival curve of TCGA-PAAD patients with high and low expression of ΔNP63 target-expressing tumors, stratified by median expression of transcripts in the PID_DELTA_NP63_PATHWAY.

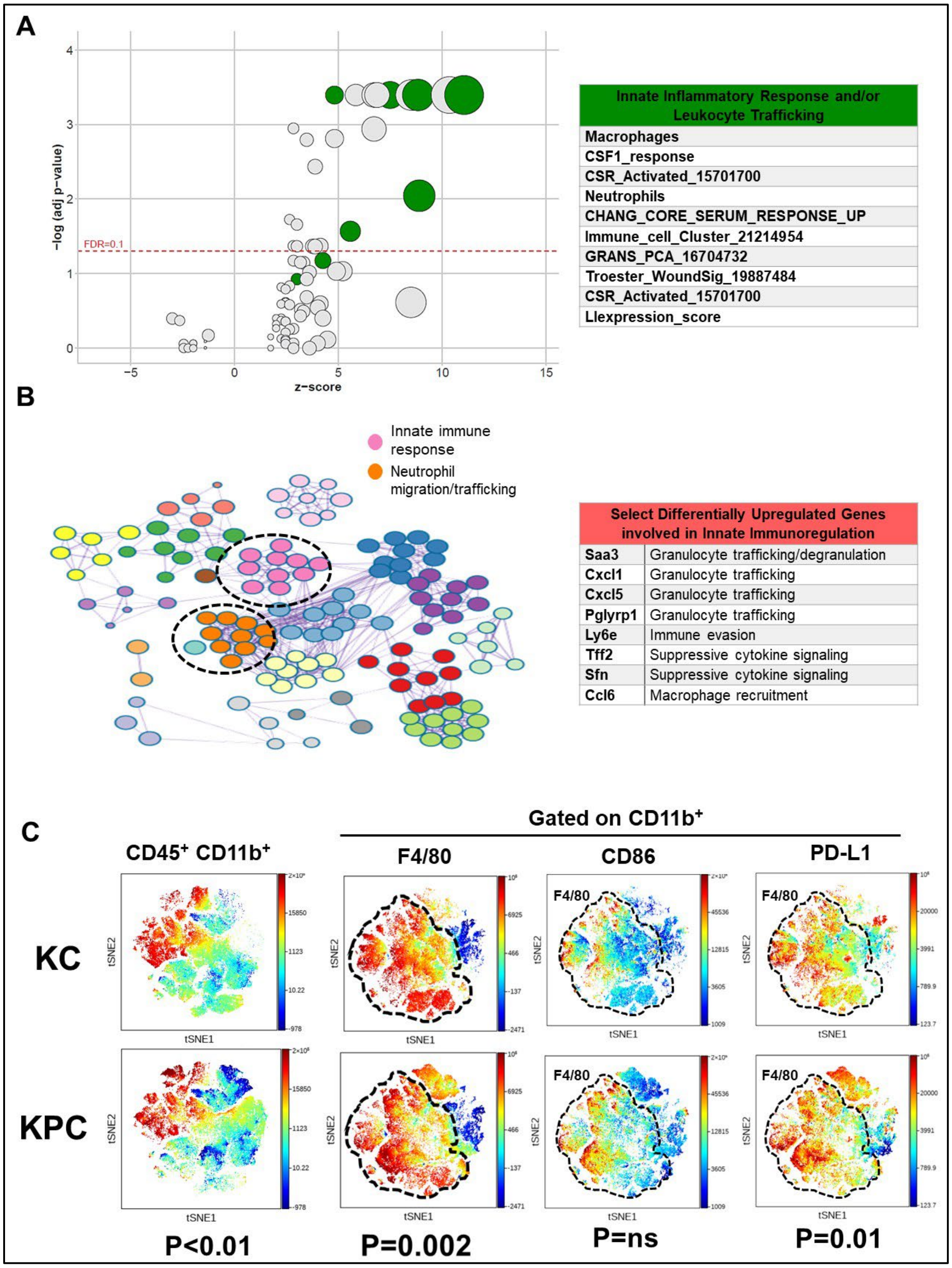


**Supplementary Figure S4**: **(A)** Bubble plot showing Pan-Immune pathways significantly differentially expressed (FDR-p<0.1) in *KRAS-TP53* co-altered vs. *KRAS*-altered/$TP53^{WT}$ TCGA-PAAD transcriptomes. Pathways specifically related to innate inflammatory response and/or leukocyte trafficking are designated in green and listed in the adjoining table; **(B)** Cytoscape network analysis depicting major nodes encoding genes involved in innate immune and neutrophil migration/trafficking in the KPC vs. KIC tumor-cell cluster via single cell RNA sequencing; **(C)** heatmaps of $CD45^{+}CD11b^{+}$ myeloid cells, as well as parent $F4/80^{+}$ and $CD86^{+}$ or $PD\text{-}L1^{+}$ expressing $F4/80^{+}$ macrophage populations in KPC and KC orthotopic tumors (n=10 mice/group) samples, overlaid in viSNE plots; dotted lines highlight the cell subset of interest.

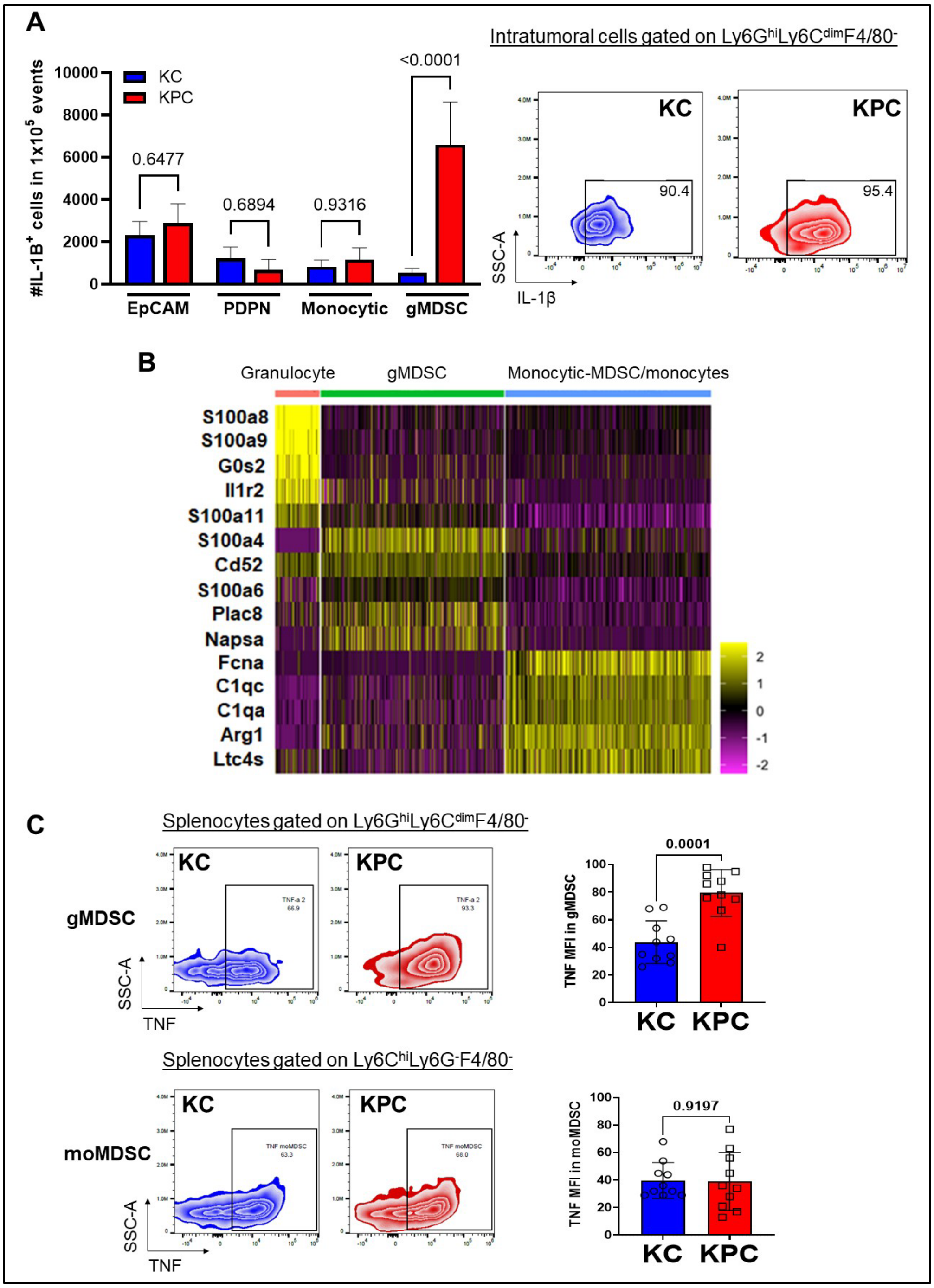


**Supplementary Figure S5**: **(A)** Histograms showing absolute number of IL-1β⁺ cells in intratumoral EpCAM⁺ tumor cells, PDPN⁺ CAFs, Ly6C⁺ monocytic cells, and F4/80⁻Ly6G⁺ gMDSC populations per $1x10^5$ live cells derived from KC and KPC orthotopic tumors (n=10 mice/group). Representative contour plot compares frequency and expression density of intracellular IL-1β in $Ly6G^{hi}Ly6C^{lo}F4/80^-$ gMDSCs from KC and KPC tumors; **(B)** Heatmap of the top-five significantly overexpressed genes defining granulocytic, gMDSC, and monocytic MDSC/monocyte sub-clusters in the combined KPC-KIC scRNAseq dataset; **(C)** Representative contour plots showing relative frequency and density of TNF expression in gMDSC ($Ly6G^{hi}Ly6C^{lo}F4/80^-$) and monocytic-MDSC ($Ly6C^+Ly6C^-F4/80^-$) in splenocytes derived from KC and KPC orthotopic tumor-bearing mice (n=10 mice/group), with adjacent histograms showing quantification.

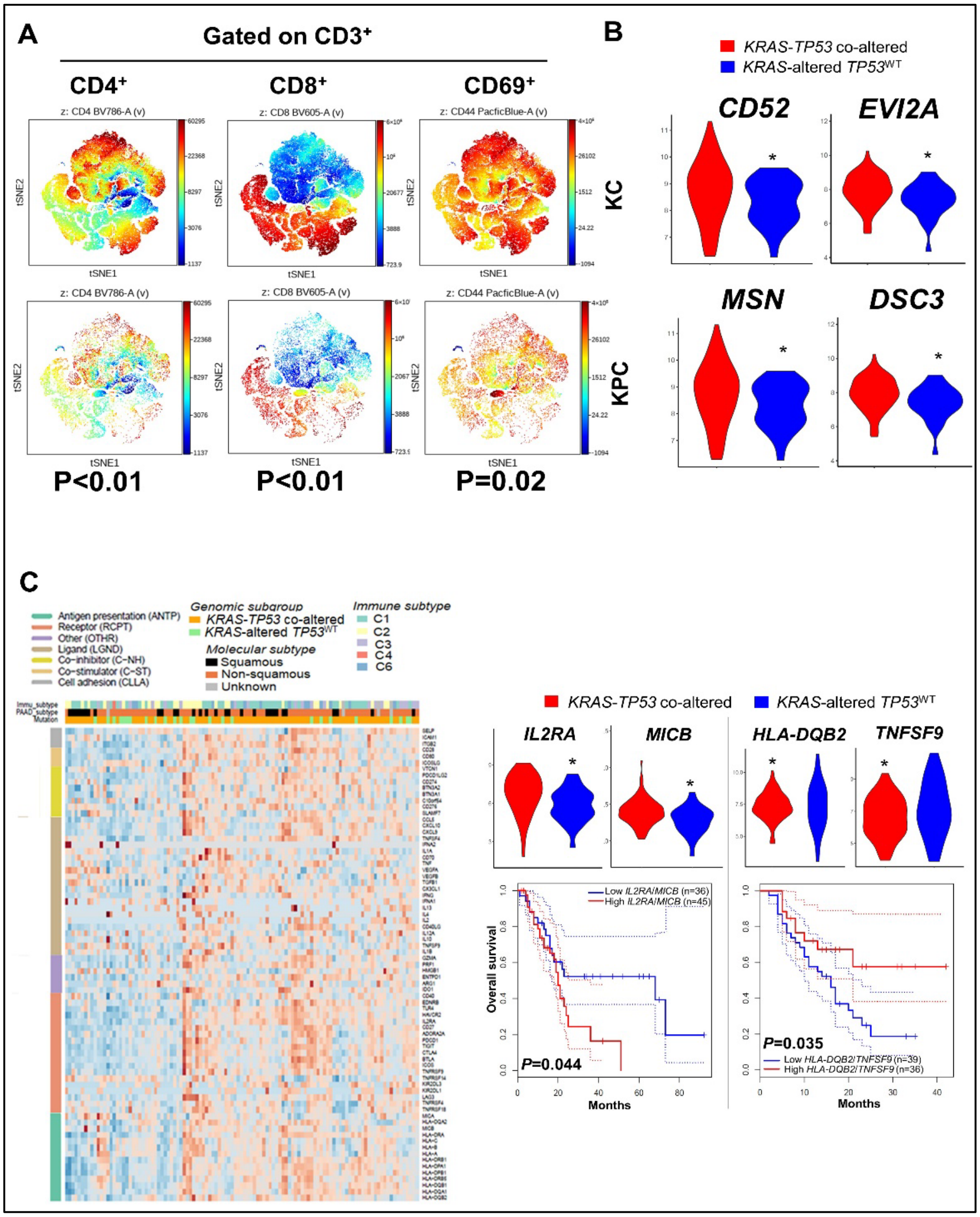


**Supplementary Figure S6: (A)** Heatmaps of CD3$^{+}$CD4$^{+}$, CD3$^{+}$CD8$^{+}$, and CD3$^{+}$CD44$^{+}$ T-cell populations in KPC and KC orthotopic tumors (n=10 mice/group) samples, overlaid in viSNE plots. Dotted lines highlight the cell subset of interest; **(B)** Violin plots showing differential expression of select genes involved in adaptive immune regulation in *KRAS-TP53* co-altered vs. *KRAS*-altered *TP53*$^{WT}$ TCGA-PAAD samples; **(C)** Heatmap representing unsupervised hierarchical clustering of ImmunoModulator-78 transcripts in TCGA-PAAD tumor transcriptomes. The heatmap is annotated by genomic subgroup (*KRAS-TP53* co-altered, *KRAS*-altered/*TP53*$^{WT}$), molecular subtype (squamous, non-squamous, unknown), and immune subtype (C1-C6); **(D)** From the ImmunoModulator-78 analysis, 4 transcripts emerged as differentially expressed in *KRAS-TP53* co-altered vs. *KRAS*-altered/*TP53*$^{WT}$ TCGA-PAAD—IL2RA, MICB, HLA-DQB2, TNFSF9, represented by violin plots. Kaplan-Meier overall survival curves in TCGA-PAAD patients with stratified by high (uppermost quartile) vs. low (lowest quartile) IL2RA/MICB and HLA-DQB2/TNFS9 expression.

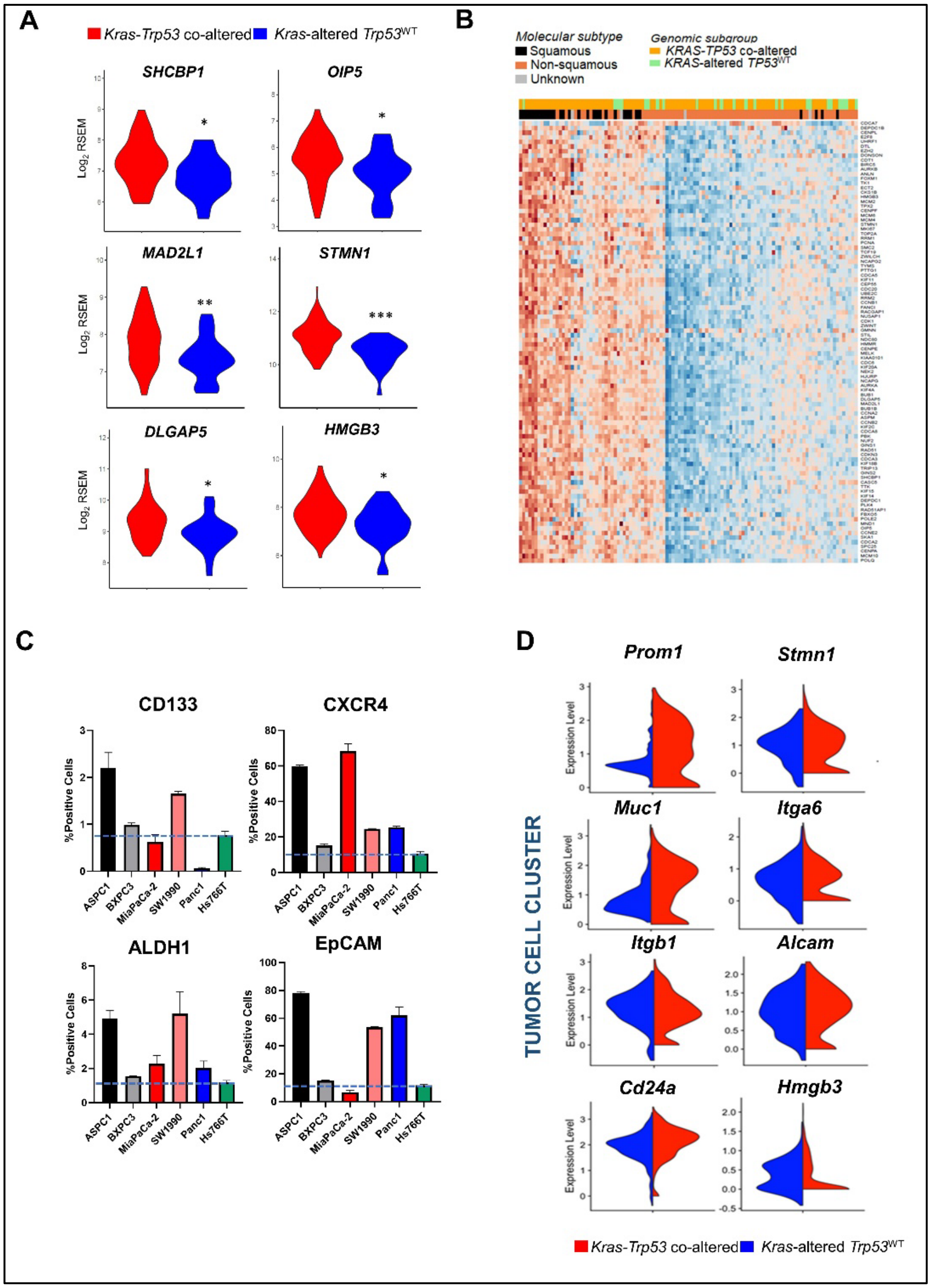


**Supplementary Figure S7**: **(A)** Violin plots showing differential expression of stemness-associated genes in the *ImSig* proliferation gene set differentially overexpressed in TCGA-PAAD *KRAS-TP53* co-altered vs. *KRAS*-altered/*TP53*$^{WT}$ samples; **(B)** Heatmap representing unsupervised hierarchical clustering of *ImSig* proliferation gene set transcripts in TCGA-PAAD tumor transcriptomes. The heatmap is annotated by genomic subgroup (*KRAS-TP53* co-altered or *KRAS*-altered/*TP53*$^{WT}$) and molecular subtype (squamous, non-squamous, unknown) in the TCGA-PAAD dataset; **(C)** Histograms showing expression of CD133$^+$, CXCR4$^+$, ALDH1$^+$, and EpCAM$^+$ in five *KRAS-TP53* co-altered (ASPC1, BxPC3, MiaPaCa-2, SW1990, and Panc1) vs. *KRAS*-altered/*TP53*$^{WT}$ (Hs766T) human PDAC cell lines via flow cytometric analysis; **(D)** Violin plots showing differential gene expression of stemness-associated genes in the tumor-cell cluster of KPC vs. KIC scRNAseq dataset.

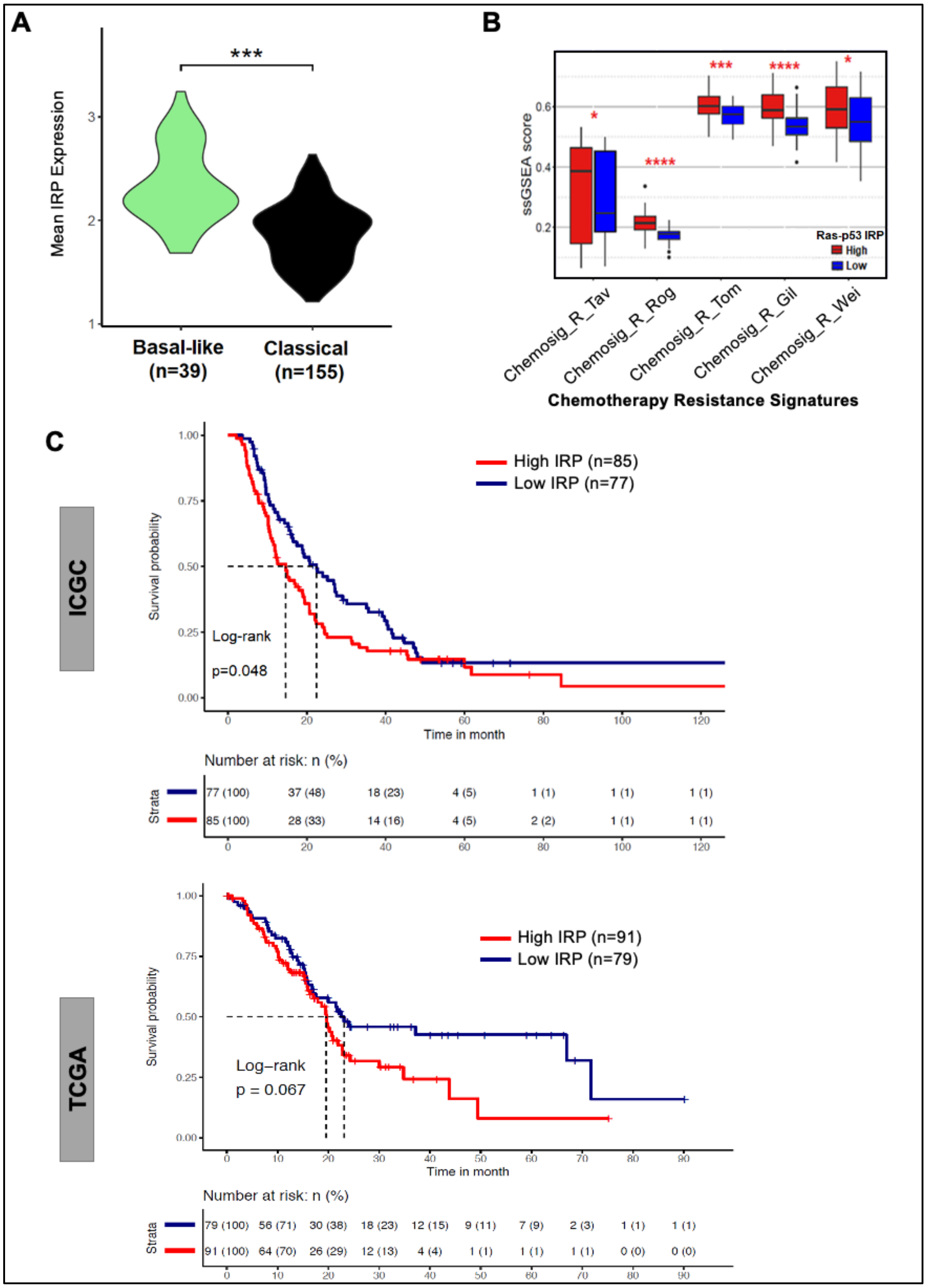


**Supplementary Figure S8**: **(A)** Violin plot showing mean *KRAS-TP53* IRP gene signature expression in transcriptomes of advanced PDAC tumor samples in the COMPASS trial, stratified into basal-like vs. classical molecular subtype; **(B)** Box and whisker plots depicting relative ssGSEA scores in *KRAS-TP53* IRP high vs. IRP low samples in five independent gene sets associated with chemoresistance in other solid tumors. Overall survival in PDAC patients, stratified by high and low expression of the *KRAS-TP53* IRP gene signature in the **(C)** ICGC and **(D)** TCGA datasets. Number at risk at each timepoint indicated in adjoining risk table.

### D. Supplementary Tables and Legends

**Supplementary Table S1**: Clinical and sociodemographic characteristics of UMiami clinical cohort of patients with advanced pancreatic cancer, stratified by *KRAS-TP53* co-altered (n=171), *KRAS*-altered *TP53*$^{WT}$ (n=42) and *TP53*-altered *KRAS*$^{WT}$ (n=32) genomic status. Separate tabs include Cox multivariable regression showing independent prognostic role of Ras-p53 co-altered status, and OncoKB annotated putative driver genes in individual patients included in the UMiami cohort.

**Supplementary Table S2**: Differentially expressed pathways in *KRAS-TP53* co-altered compared to *KRAS*-altered *TP53*$^{WT}$ TCGA-PAAD samples, categorized into MSigDB:C2CP v6.2 ("canonical pathways") and Pan-Immune signature sets.

**Supplementary Table S3**: Pathways related to stemness derived from studies examining stemness properties in cancer and/or embryonic cells.

**Supplementary Table S4**: Tabulation of Cancer Cell Line Encyclopedia (CCLE) human pancreatic ductal adenocarcinoma (PDAC) cell lines (n=41) stratified by K*RAS-TP53* genomic status, Bailey molecular subtype classification, normalized expression of gene transcripts defining squamous transdifferentiation, and mRNA stemness index score.

**Supplementary Table S5**: Immune-related gene set enrichment analysis (GSEA) used for immune deconvolution.

These files can be found at: https://doi.org/10.6084/m9.figshare.19689256

E. **Supplementary Table S6**: Twenty genes differentially overexpressed in *KRAS-TP53* co-altered PDAC—associated with squamous transdifferentiation, innate immunoregulation, adaptive immune evasion, inflammasome machinery, and stemness features—defined as *KRAS-TP53* immunoregulatory program (IRP).

| |
|---|
| PANX1 |
| HMGB3 |
| IL18 |
| THBD |
| CD163 |
| FCGR2A |
| DLGAP5 |
| STMN1 |
| SCPEP1 |
| SHCBP1 |
| OIP5 |
| MAD2L1 |
| MICB |
| LY96 |
| AQP9 |
| EVI2A |
| IL2RA |
| CD300 |
| CD52 |
| NAMPT |

F. **Supplementary Table S7**: List of antibodies used for flow cytometry and histologic/western blotting analysis in *in-vitro* and *in-vivo* experiments.

| **FLOW CYTOMETRY** | | | | |
|---|---|---|---|---|
| **PRIMARY ANTIBODY** | **FLUOROPHORE** | **SUPPLIER** | **SPECIES** | **CATALOGUE NUMBER** |
| **CD45** | FITC | Biolegend | Mouse | 103108 |
| **CD45** | BV510 | Biolegend | Mouse | 103137 |
| **CD3** | PerCPCy5.5 | Biolegend | Mouse | 100218 |
| **CD4** | BV785 | Biolegend | Mouse | 100453 |
| **CD8** | BV605 | Biolegend | Mouse | 100744 |
| **CD44** | APC-Cy7 | Biolegend | Mouse | 103028 |
| **CD62L** | PE | Biolegend | Mouse | 104408 |
| **PD-1** | PE-Cy7 | Biolegend | Mouse | 135216 |
| **CD107a** | PE-Dazzle | Biolegend | Mouse | 121624 |
| **CD11b** | AF700 | eBioscience | Mouse | 56-0112-82 |
| **Cd11c** | BV421 | Biolegend | Mouse | 117343 |
| **F4/80** | BV785 | Biolegend | Mouse | 123141 |
| **Ly6G** | BV650 | Biolegend | Mouse | 127641 |
| **Ly6G** | BUV563 | BD | Mouse | 612921 |
| **Ly6C** | PerCPCy5.5 | Biolegend | Mouse | 128012 |
| **Ly6C** | APC-Cy7 | Biolegend | Mouse | 128026 |
| **CD206** | PE-Cy7 | Biolegend | Mouse | 141720 |
| **CXCR2** | FITC | Biolegend | Mouse | 149310 |
| **PDPN** | PerCPCy5.5 | Biolegend | Mouse | 127422 |
| **EpCAM** | BV605 | Biolegend | Mouse | 118227 |
| **Il1-beta** | PECy7 | Invitrogen | Mouse | 25-7114-82 |
| **TNF-alpha** | PE | Biolegend | Mouse | 506306 |
| **CD133** | PE-eFluor610 | eBioscience | Human | 61-1338-42 |
| **CXCR4** | PE-Cy7 | eBioscience | Human | 25-9999-42 |
| **ALDH1** | - | ThermoFisher | Human | PA5-11537 |
| **EpCAM** | AlexaFlour488 | Biolegend | Human | 324210 |
| **L/D** | Blue | Invitrogen | | L34962A |

| **HISTOLOGY/WESTERN BLOT** | | | | |
|---|---|---|---|---|
| **PRIMARY ANTIBODY** | **CLONE** | **SUPPLIER** | **HOST SPECIES** | **CATALOG NUMBER** |
| **Ly6G** | RB6-8C5 | Abcam | Rat | ab25377 |
| **F4/80** | D2S9R | Cell Signaling | Rabbit | 70076 |
| **DeltaNp63** | EPR17863-47 | Abcam | Rabbit | ab203926 |
| **CD8** | D4W2Z | Cell Signaling | Rabbit | 98941 |
| **Vinculin** | E1E9V | Cell Signaling | Rabbit | 13901 |
| **CD133** | RM1002 | Abcam | Rabbit | ab19898 |
